# Hippocampal timestamp for goals

**DOI:** 10.1101/2023.07.27.550892

**Authors:** Alison Montagrin, Denise E. Croote, Maria Giulia Preti, Liron Lerman, Mark G. Baxter, Daniela Schiller

**Author notes:** ***Corresponding Authors:*** Alison Montagrin, University of Geneva, Geneva, 1202, Geneva, Daniela Schiller, Icahn School of Medicine at Mount Sinai, New York, NY 10029. these authors contributed equally (Alison Montagrin, Denise E. Croote).

## Abstract

Our brain must manage multiple goals that differ in their temporal proximity. Some goals require immediate attention, while others have already been accomplished, or will be relevant later in time. Here, we examined how the hippocampus represents the temporal distance to different goals using a novel space-themed paradigm during 7T functional MRI (n=31). The hippocampus has an established role in mental time travel and a system in place to stratify information along its longitudinal axis on the basis of representational granularity. Previous work has documented a functional transformation from fine-grained, detail rich representations in the posterior hippocampus to coarse, gist-like representations in the anterior hippocampus. We tested whether the hippocampus uses this long axis system to dissociate goals based upon their temporal distance from the present. We hypothesized that the hippocampus would distinguish goals relevant for ones’ current needs from those that are removed in time along the long axis, with temporally removed past and future goals eliciting increasingly anterior activation. We sent participants on a mission to Mars where they had to track goals that differed in when they needed to be accomplished. We observed a long-axis dissociation, where temporally removed past and future goals activated the left anterior hippocampus and current goals activated the left posterior hippocampus. Altogether, this study demonstrates that the timestamp attached to a goal is a key driver in where the goal is represented in the hippocampus. This work extends the scope of the hippocampus’ long axis system to the goal-mapping domain.

## Main

Tracking personal goals is a vital and ongoing cognitive process. Our brains are continuously monitoring what we have already accomplished, are actively undertaking, and are planning to tackle in the future. For instance, we know that we no longer need to go to the grocery store, but we do need to pay the credit card bill today to avoid accruing interest, and mail a wedding gift while in town tomorrow. Though all of these goals are occupying our mental spaces, they differ on a fundamental parameter: time. Paying the credit card bill is a goal that must be accomplished in the present. The other tasks are removed in time from the needs of the current self and represent goals that were either completed in the past or remain to be accomplished in the future. Yet, goals are dynamic entities. Mailing the wedding gift begins as a future goal, but transitions to a current priority and a task of the past as we move forward in time. Therefore, effectively tracking personal goals involves managing different goals and updating the relevancies of each goal in memory as time progresses.

Episodic memory is a system that receives and stores information about episodes that are temporally dated and linked by temporal-spatial relations (Tulving, 1984). The hippocampus has long been known to be critical in episodic memory processing (e.g., Scoville, 1957; Tulving, 1984; Tulving & Markowitsch, 1998). Episodic memories encoded in the hippocampus are not only employed to travel back in time but also to project oneself into the future (D’Argembeau et al., 2008; Tulving, 1986; Williams et al., 2007). A key role of episodic memory might be to adjust behavior to respond to current and future goals in remembering preferentially goal-relevant events (Conway et al., 2019; Montagrin and Sander, 2016; Montagrin et al., 2018; Williams et al., 2008). Studies of spatial navigation in rodents and humans have shown that a critical role of the hippocampus is to locate relevant goals in the environment, and to encode or remember temporal or/and spatial information of events (e.g., food; Chadwick et al., 2015; Eichenbaum, 2014; Fortin et al., 2002; Howard et al., 2014; Kesner et al., 1988; Mayes et al., 2001; Nyberg et al., 2021; O’Keefe and Nadel, 1978; Olafsdottir et al., 2019; Palombo et al., 2017; Shimamura et al., 1990). But how does the brain keep an updated representation of goals that have been accomplished versus goals that require immediate attention or goals that will be relevant later in time?

We investigated how the timestamp attached to a goal is processed and represented in the human brain using functional magnetic resonance imaging (fMRI). We focused our investigation on the hippocampus, due to its established role in memory and mental time travel (Eichenbaum, 2000; Schacter et al., 2007; Tulving & Markowitsch, 1998), and examined whether the hippocampus maintains an updated representation of goals’ temporal proximity. Substantial literature has identified anatomical and functional specialization along the hippocampal longitudinal axis, notably in the episodic and spatial domains (Poppenk et al., 2013; Robin & Moscovitch, 2017; Strange et al., 2014). Here, we asked whether goals are also mapped along the hippocampal long-axis based upon their temporal distance from the present.

In our paradigm, participants embarked on an imaginary 4-year mission to Mars where they needed to complete a series of goals in order to survive on the planet. These goals varied in when they needed to be accomplished during the mission, with different sets of goals applicable in the first, second, third, and fourth year of the trip. As participants advanced through the mission, they were shown the *same* goals, but they themselves moved forward in time, which shifted the goals’ temporal relevancies. Future goals transitioned to current needs, while current needs became tasks of the past. Thus, participants had to manage multiple goals that differed in their temporal distance and update their goal representations as they moved through their 4 years on Mars. We hypothesized that current goals would activate more posteriorly in the hippocampus, due to their temporal proximity to the present, and temporally removed goals would activate more anteriorly, due to their temporal distance from the present. To focus our analyses on the hippocampal long axis, we combined multi-echo multi-band imaging with ultra-high field 7T fMRI (Kundu et al., 2012, 2017).

We framed the goals in the context of a space mission using fixed goal properties (i.e., their names and visual features), rather than employing goals from participants’ personal pasts and futures, in order to isolate the temporal distance parameter. When comparing representations of items that are close in time to those that are far in time, be it past memories or future projections, there are inherent differences in the content, realness, and level of detail associated with the constructions (Gilmore et al., 2021; Liberman & Trope, 2014; Trope & Liberman, 2010). These experiential differences can then cloud the interpretation of consequent behavioral and neural observations. Our paradigm addressed this limitation by holding the properties of the goals fixed, and the level of detail associated with the present, past, future, near, and distant conditions constant. Thus, the only thing that changed about a goal throughout the experiment was its temporal distance from the present.

## Results

Thirty-four neurologically healthy individuals completed the two-day experiment and 31 remained in the sample after MRI exclusions (*M*=27.0 years, *SD*=4.5, Range=19-36, n=15 females; see Methods). Participants were taught when they needed to complete each goal via a training exercise on the first day of the experiment (**Figure 1a**). In addition to learning goals relevant for the first, second, third, and fourth years of the mission, participants also learned a set of goals that they always needed to accomplish and a set of goals that they never needed to attend to. Participants learned the relevancy of five goals for each of the 6 aforementioned timeframes, totalling to 30 stimuli (see Methods for training details).

**Figure 1:**
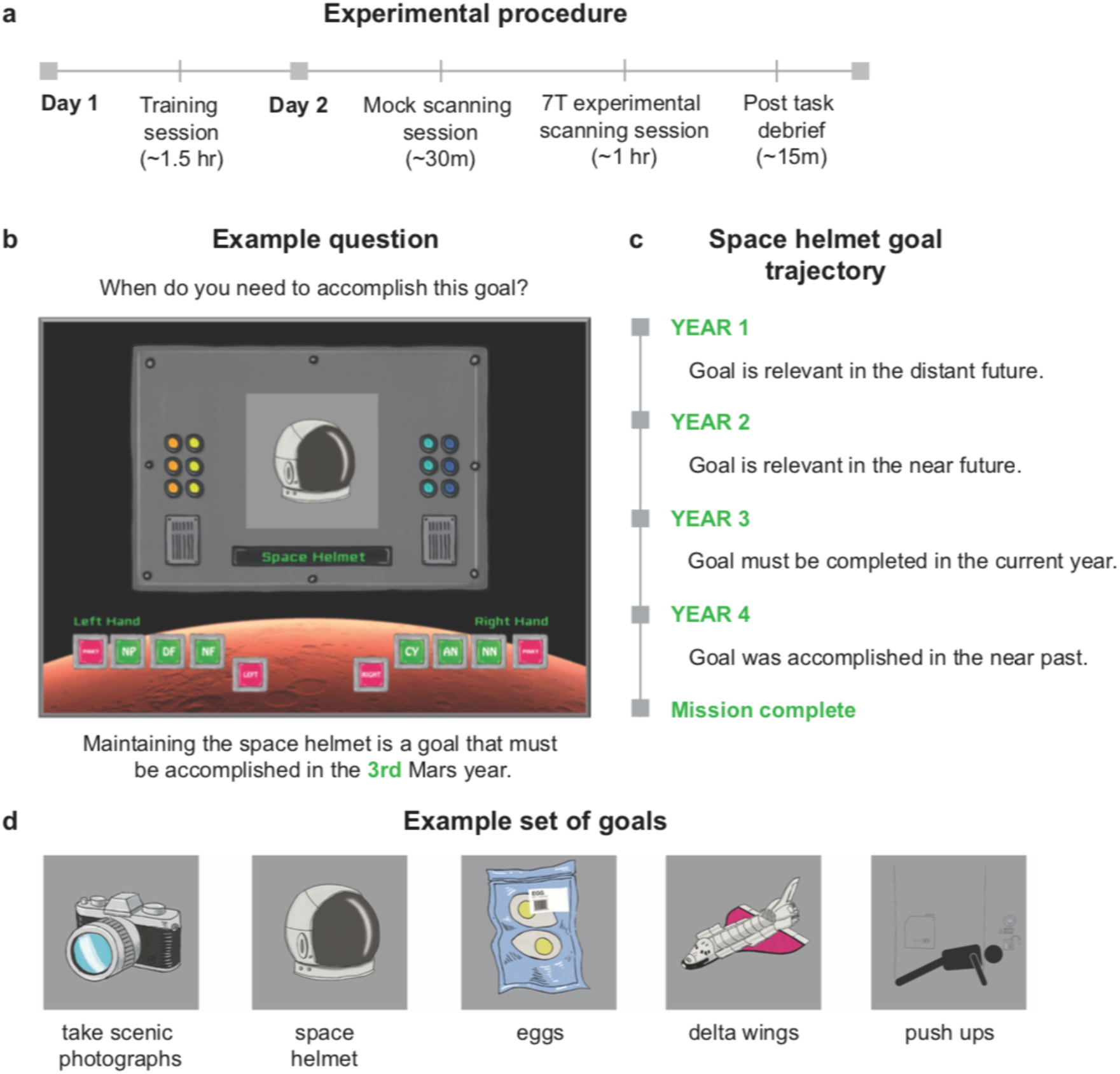
Experimental design. **(a)** The experiment took place over two days. Participants learned when they would need to complete each goal during a training session on the first day of the experiment. Participants were familiarized with the paradigm in the mock scanner and then embarked on their mission inside the 7T scanner on the second day. **(b)** Example screen shown during year 1. Participants were shown a goal and asked when they need to accomplish this goal, with buttons at the bottom of the screen to indicate whether the goal was applicable for the current Mars year (CY), or relevant in the distant future (DF), near future (NF), near past (NP), distant past (DP), always (AN, always needed), or never (NN, never needed). **(c)** Example trajectory of the space helmet goal across the mission, which is relevant in the 3^rd^ Mars year for participants completing version 1. **(d)** Example set of goals. Participants learned 6 sets of 5 goals, each set associated with either year 1, year 2, year 3, year 4, always, or never.

On the second day of the experiment, participants were trained on the paradigm in a mock scanner and then sent on their mission in an ultra-high field 7T MRI scanner (**Figure 1a**). They were shown a goal and asked when they need to accomplish it, with buttons present to indicate whether the goal was applicable for the current Mars’ year, or relevant in the distant past, near past, near future, distant future, always, or never (**Figure 1b**). Distant goals were defined as those 2-3 years from the present and near goals within 1 year of the present during the mock scanner training session. Participants evaluated when they needed to accomplish each of the 30 goals before advancing to the next year of the mission. They were then shown the *same* 30 goals, but the goals’ temporal relevancies shifted now that the participants had moved forward in time (**Figure 1c**). For example, when participants were in their 2^nd^ year of the expedition and were presented with a goal relevant in the 3^rd^ year, they selected the near future button (NF) on the screen to indicate that this goal should be attended to in the near future. When they advanced to the 3^rd^ year of the game, this goal shifted to a current priority and participants selected the current year button (CY). Therefore, participants had to continuously apply their knowledge of the timing of the goals and imagine when they needed to complete each goal in relation to their own temporal position (see Methods for a full task description).

As the mission progressed, new buttons appeared on the screen allowing participants to indicate whether they had already completed the goal displayed in the past, and buttons were removed from the screen as the future timeframe disappeared throughout the mission. The trials were divided into instances where participants were making an evaluation of a goal that needed to be completed in the distant future (DF), near future (NF), near past (NP), and distant past (DP; 15 trials each), or in the current year (CY), always needed (AN), and never needed (NN; 20 trials each), totalling to 120 trials. All goals revolved around space shuttle maintenance, space suit care, personal nutrition, exercise, and recreational activities (**Figure 1d**).

## Behavioral results

### Participants took longer to process temporally removed goals than current goals

We first explored the behavioral impact of the timestamps attached to the goals. We compared reactions times for goals that were removed in time (distant future, near future, near past, distant past) to reaction times for current goals. If participants are using a mental time travel-like process to evaluate the goals, we expected longer response times for the temporally removed goals in the past and future compared to the goals in the present year.

We examined participants’ reaction times using a series of linear mixed models, implemented via the lme4 (Bates et al., 2015) and lmerTest (Kuznetsova et al., 2017) packages in R v. 3.6.0 (R Core Team, 2013). After removing incorrect and reaction time outlier trials (+/- 3 SD’s of a participant’s mean) 94.5% of all trials remained. We modeled participants’ log transformed reaction times against temporal condition, with levels for distant future, near future, current, near past, and distant past trials. Current trials were distributed equally throughout the game, while future trials were presented in the beginning 75% and past trials in the last 75% of the mission. This distribution was in effect because there were no future goals in the last year of the mission, and no goals had already been completed in the first year of the mission (see **Supplementary Table 1** for a visualization of the trial numbers).

We included a random effect for participant ID to account for the lack of independence between trials within a participant, and game year was included as a fixed effect and as a random effect for objective time to account for any trends in reaction time within and between participants across the years as the mission proceed (Table 1). This game year regressor has four levels (year 1, year 2, year 3, year 4) and was in place to absorb variance such that any reaction time differences that remained between the conditions were those above and beyond what could be explained by time point in the game.

**Table 1.**
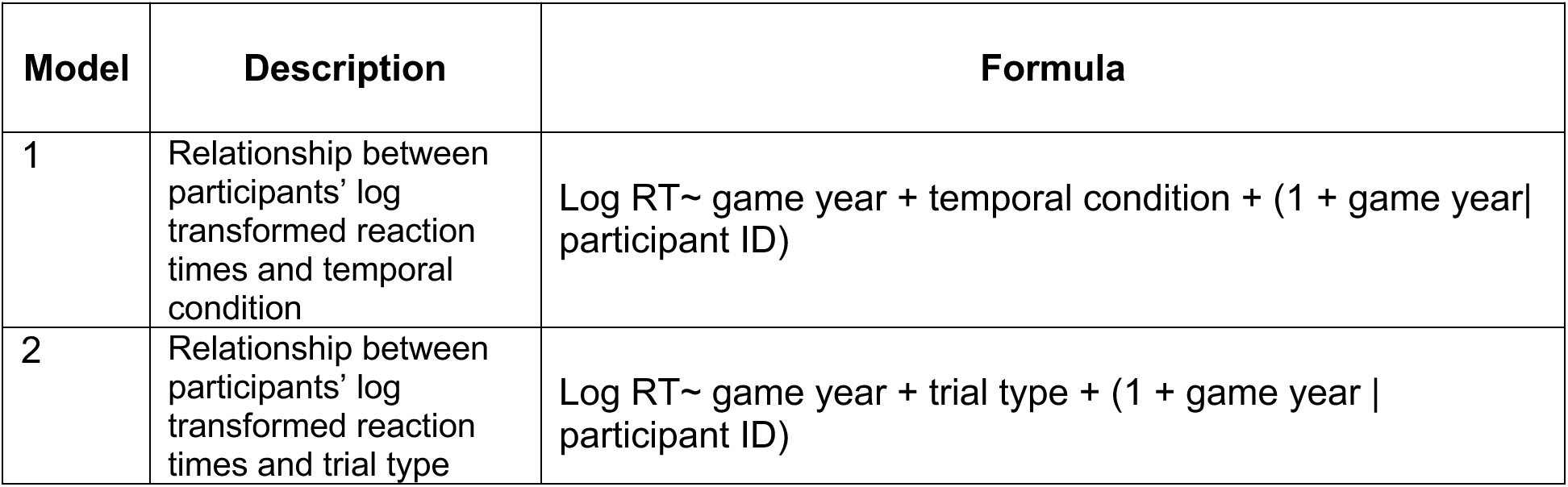
Reaction time linear mixed models. The dependent variable was comprised of participant’s log transformed reaction times for each trial. The temporal condition predictor in Model 1 included five levels, distant future, near future, near past, distant past, and current trials. Always and never trials were filtered from the dataset for Model 1. The trial type predictor in Model 2 included two levels, trials with a temporal element (distant future, near future, near past, distant past trials) and trials without (current, always, never trials). The participant ID random effect contained 31 participants and was included to account for the lack of independence between trials within a participant. The game year fixed effect and random effect included 4 levels (year 1, year 2, year 3, year 4) and served to account for downward reaction time trends as the game progressed between and within participants.

We evaluated statistical significance using Type II Sums of Squares and computed the model-based estimated marginal means, standard errors, and confidence limits via the emmeans package in R. All *post hoc* contrasts quantified the difference between the two estimated marginal means and outputted the corresponding estimate, standard error, test statistic, and *p* value (Lenth et al., 2021). *Post hoc* tests were corrected for multiple comparisons using a Tukey adjustment.

The reaction times to process the same goals shifted based upon the trajectories of the goals, i.e., how their temporal relevance changed across the game (**Figure 2a**). This suggests that participants were indeed reframing how they thought about each goal as its temporal context shifted. Participants’ log-transformed reaction times were significantly associated with temporal condition (Type II ANOVA; *F*(6, 3416) = 87.07, *p*<0.0001). *Post hoc* comparisons revealed that, as hypothesized, current trials were processed faster than all temporally removed conditions when contrasted individually (**Figure 2b, c; Supplementary Table 2**).

**Figure 2:**
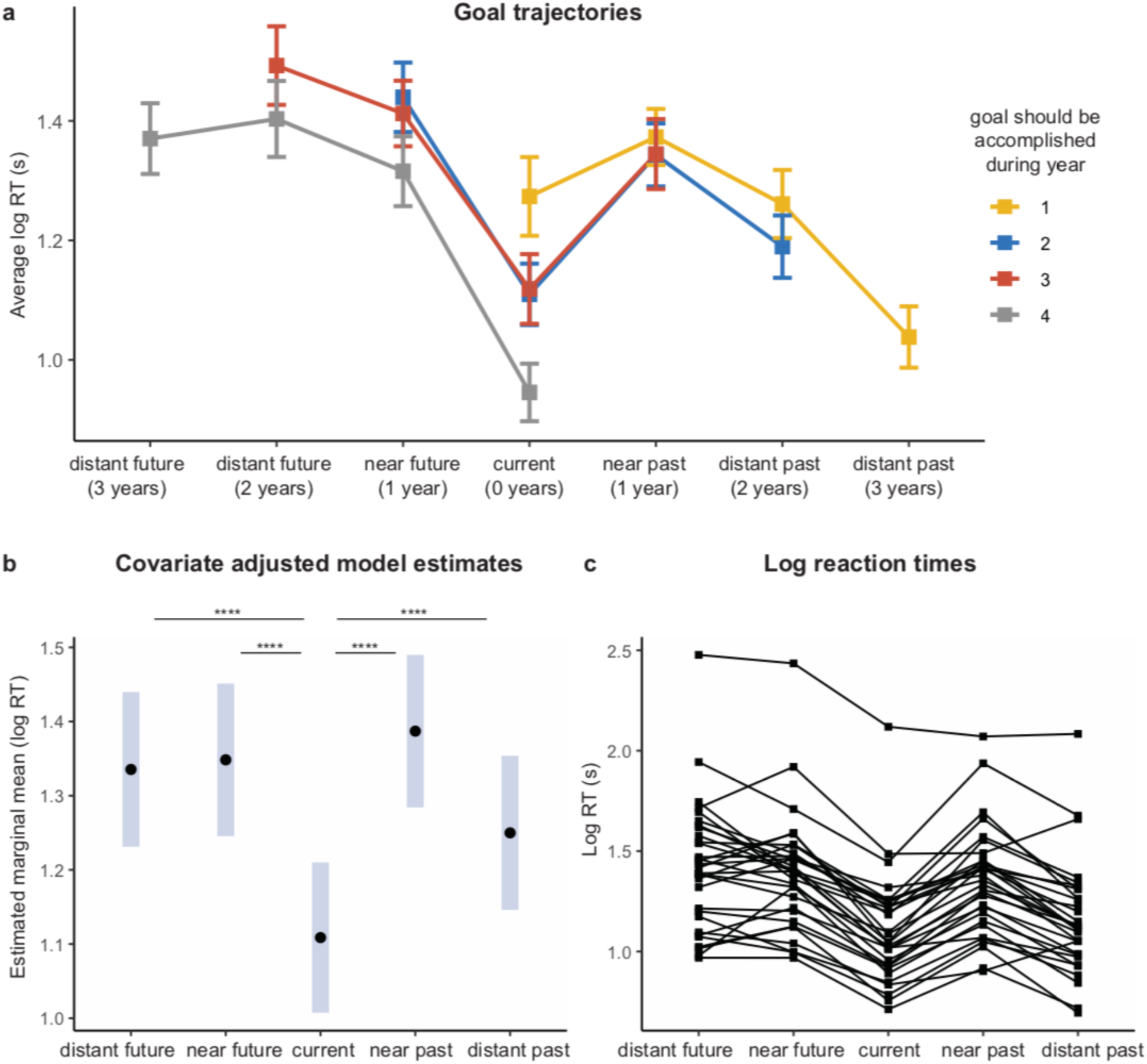
Participants took longer to process temporally removed goals than current goals. **(a)** Participants shifted their thinking of the goals as the goals’ temporal relevance changed. This plot visualizes the trajectories of the goals, separated by when they had to be accomplished in the game. Error bars = standard error of the mean (SEM). **(b)** *Post hoc* comparisons of the estimated marginal means revealed that current goals were processed faster than all temporally removed goals (**Supplementary Table 2**). Error bars = 95% CI. **(c)** Individual participants’ average log reaction time for each of the temporal conditions. **** *p*<0.0001.

We validated the findings above using a second model that also included the always and never trials. As with the current goals, always and never goals did not include a temporal element. We separated trials into those involving a temporal computation (distant future, near future, near past, distant past) and those not (current, always, never) and likewise found that participants took significantly longer to process trials involving a temporal element (**Supplementary Figure 1**). In summary, our behavioral findings provide compelling evidence that the brain evaluates goals relevant to one’s current situation differently than it does those that are removed in time.

In an additional type of trials, participants had to choose between two goals: one relevant for the current year and one relevant for another timeframe. We used a similar linear mixed model as above to examine the additional choice trials (see **Supplementary Figure 2**). *Post hoc* comparisons of the estimated marginal means revealed that when participants had to choose between current and distant future goals, they processed faster than when participants had to choose between current and near future or between current and near past. These results possibly reflect the pattern separation theory (e.g., Lohnas et al., 2018). The boundary between the goals in the current moment and those that are near in time is more blurred compared to goals that are further away in time and where the boundary with the present would be sharper and would take less time to distinguish.

## Neuroimaging results

### Timestamp for goals represented along the longitudinal axis of the hippocampus

We sought to identify regions within the hippocampus that showed differential activity for temporally removed and current goals. We defined a series of general linear models (GLMs) with regressors for each temporal condition, the always and never trials, and years 1 – 4 of the game (see Methods for full model descriptions and **Supplementary Tables 10 and 11**). We first contrasted blood-oxygen-level-dependent (BOLD) activity for the grouped temporally removed conditions (distant future, near future, distant past and near past) with that of the current condition, and vice versa, bilaterally in the hippocampus (see **Supplementary Figure 3** for the parameter estimates). We used an automated subcortical segmentation methods FreeSurfer v7.2 to measure volumes of these regions of interest (i.e., bilateral hippocampus, see Methods for more details).

We hypothesized that signal amplitude would differ along the long axis as a function of temporal distance. Consistent with this hypothesis, temporally removed (Remote condition) goals produced stronger activation than current (Current condition) goals in the left anterior hippocampus (**Figure 3a**; **Table 2**). The contrast Remote versus Current identified voxels falling anterior to Y=-21 (mm in MNI coordinate). Current goals produced stronger activation in the left posterior hippocampus, with voxels extending from Y=-31 to Y=-36 (**Figure 3b**; **Table 2**). Thus, the *same* goals were anatomically dissociated along the longitudinal axis based on whether they were currently relevant, or relevant at a point removed in time (**Figure 3c, d**).

**Figure 3:**
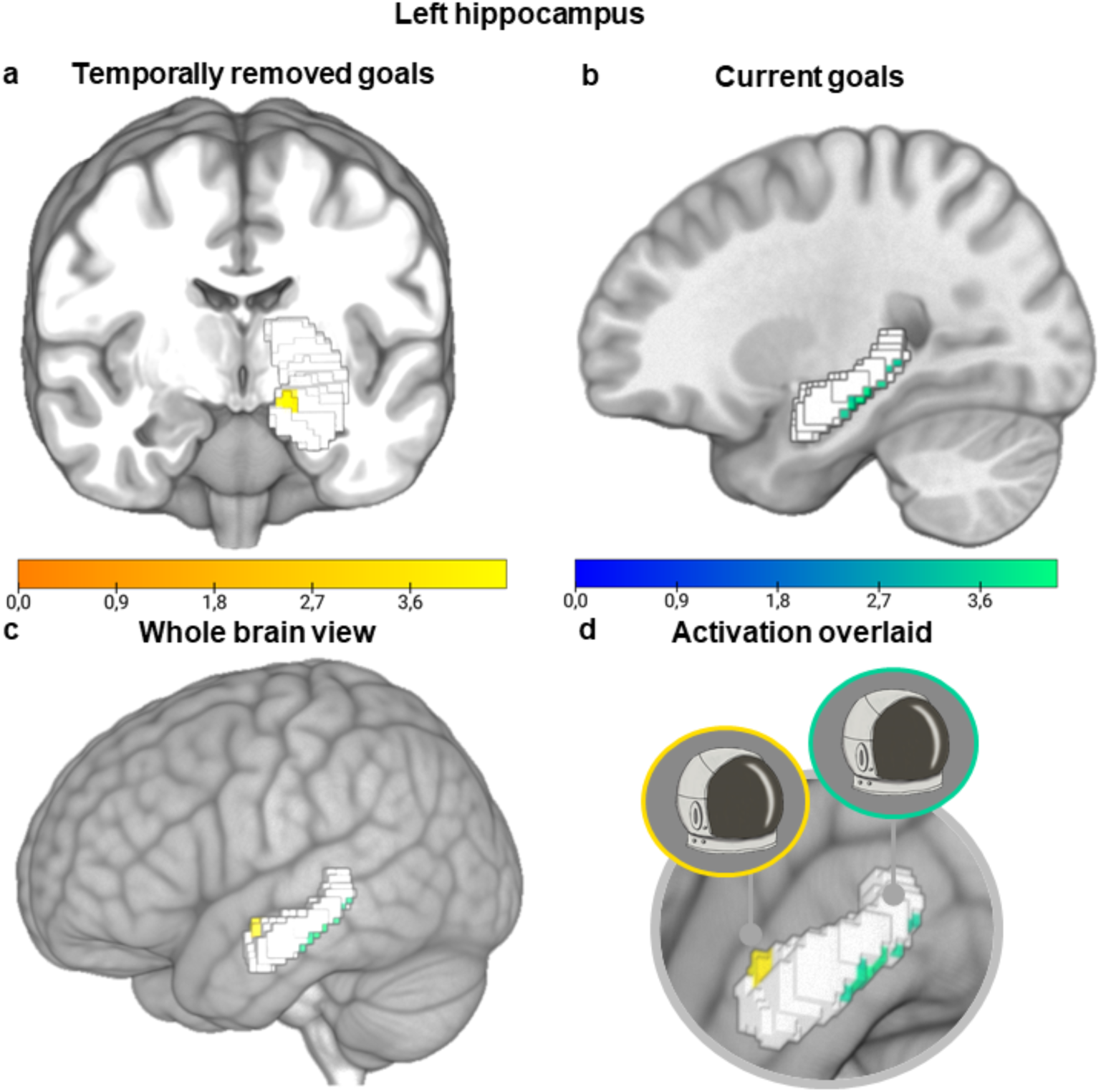
Temporally removed goals activated the left anterior hippocampus and current goals activated the left posterior hippocampus. **(a)** Activation maps for the contrasts comparing the remote (distant future + near future + distant past + near past) > current are overlaid in yellow. **(b)** Activation maps for the contrasts comparing the current > remote are overlaid in green. All z-statistic images were thresholded parametrically using GRF-theory-based maximum height thresholding with a (FWE-corrected) significance threshold of *p*=0.025. **(c)** Activation for the temporally removed goals (yellow) and the current goals (green) shown concurrently on the brain. **(d)** The *same* goal, for instance fixing the space helmet, was anatomically dissociated along the longitudinal axis based on whether it was currently relevant, or relevant at a point removed in time. Contrast maps were overlaid and rendered onto a 3-dimensional MNI 152 brain using MRIcroGL. The left hippocampal ROI is displayed in light gray. Color bars reflect the thresholded z-statistic scores.

**Table 2:**
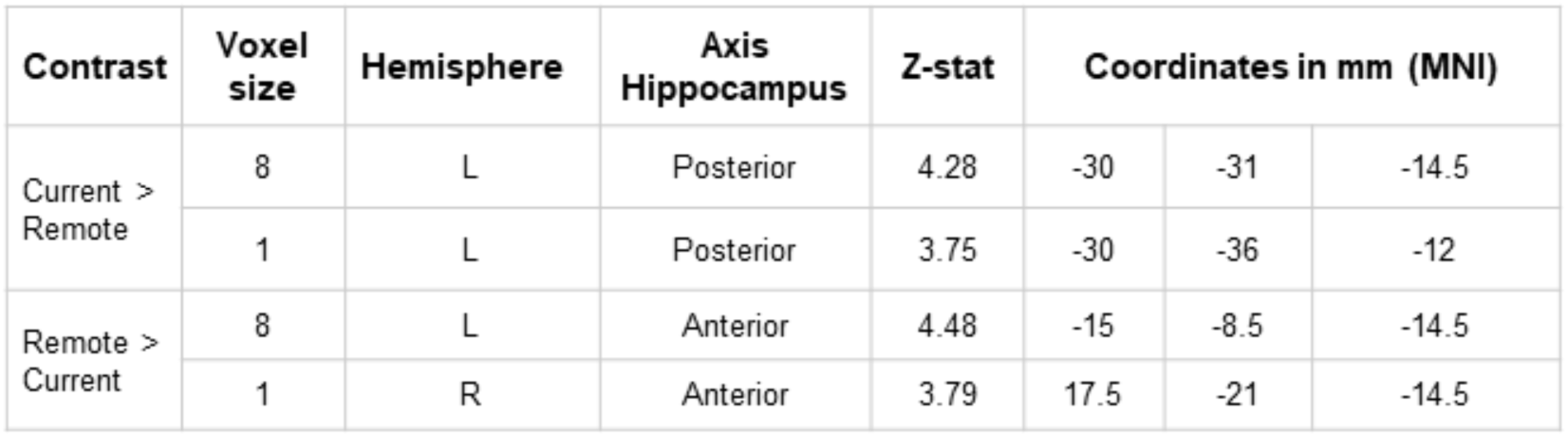
Coordinates of peak activation for temporally removed and current goals in the left and right hippocampus. Temporally removed (past and future) goals activated the left anterior hippocampus, while current goals activated mainly the left posterior hippocampus, and one voxel in the right hippocampus. This table reports the number of significant voxels in the cluster, maximum z-statistic within the cluster, and the x, y, z location of the maximum intensity voxel for each contrast. Coordinates are reported in Montreal Neurological Institute (MNI) space. All statistical maps were corrected for multiple comparisons using maximum height thresholding (FWE voxel-wise correction, *p*=0.025).

| Contrast | Voxel size | Hemisphere | Axis Hippocampus | Z-stat | Coordinates in mm (MNI) |  |  |
| --- | --- | --- | --- | --- | --- | --- | --- |
| Current > Remote | 8 | L | Posterior | 4.28 | -30 | -31 | -14.5 |
|  | 1 | L | Posterior | 3.75 | -30 | -36 | -12 |
| Remote > Current | 8 | L | Anterior | 4.48 | -15 | -8.5 | -14.5 |
|  | 1 | R | Anterior | 3.79 | 17.5 | -21 | -14.5 |

As a secondary hypothesis, we examined whether at the brain level, the hippocampus could account for a temporal gradient along its axis, although the behavioral results did not show a difference in a temporal gradient. We contrasted BOLD activity for each temporally removed past and future conditions with that of the current condition and vice versa specifically in the left hippocampus as the previous results were mainly in the left hemisphere (see **Supplementary Table 3**).

All analyses were corrected for multiple comparisons implementing Family-Wise Error (FWE) using GRF-theory based maximum thresholding (voxel-wise correction, two-tailed *p*=0.025). These voxel locations align with the posterior, and anterior boundaries outlined in Poppenk et al. (2013), who defined the foci of the anterior hippocampus to y = -21, using the uncal apex as the anatomic landmark. Although we observed this strong long axis distinction in the left hippocampus, activation differences were not as prominent in the right hippocampus (**Table 2**).

Though our a priori hypotheses were centered on the hippocampus, we further conducted an exploratory whole brain analysis. This GLM was designed as described above. All statistical maps were masked with a gray matter tissue probability map and corrected for multiple comparisons using voxel-wise correction thresholding (FWE voxel-wise correction, *p*=0.025). Temporally removed goals largely activated anterior brain region, whereas current goals more strongly activated posterior brain regions (**Supplementary Figure 4, Table 4**). This separation, namely the localization of temporally removed goals to anterior brain regions, is in line with the role of the frontal cortex in autobiographical planning and future mental time travel (Ciaramelli et al., 2021; Spreng et al., 2010). An exploratory analysis of functional connectivity is consistent with this hypothesis (see **Supplementary Figure 5**).

In summary, the left hippocampus anatomically distinguished present goals from goals that were removed in time along its longitudinal axis. Temporally removed past and future goals activated the anterior hippocampus, while current goals activated a more posterior portion of the hippocampus.

## Discussion

In this study we investigated whether goals are mapped along the hippocampal anterior-posterior axis based on when they need to be accomplished. We examined how participants represented goals that were a current priority, goals that they accomplished in the past, and goals that they needed to complete in the future using a space mission themed paradigm. Behaviourally, goals that were relevant for ones’ current needs were processed faster than temporally removed past and future goals. On a neural level, current goals activated the left posterior hippocampus and temporally removed past and future goals activated the left anterior hippocampus. Altogether, these findings extend the known scope of the hippocampus’s long-axis system to the goal-mapping domain. They demonstrate that the mental timestamps assigned to goals that are otherwise identical, guide their dissociation along the anterior-posterior parts of the hippocampus.

Long axis specialization has been explored in the episodic memory and spatial navigation domains. This works has largely focused on differences in representational granularity along the longitudinal axis and has found evidence of a detail-rich to gist-like gradient system (Poppenk et al., 2013; Robin & Moscovitch, 2017; Strange et al., 2014). Small scale event information (Collin et al., 2015) and detailed spatial information (Nadel et al., 2013) are represented in the posterior hippocampus, while multi-level narrative information (Collin et al., 2015) and contextual spatial information (Nadel et al., 2013) activate the anterior hippocampus. These functional divisions are supported by anatomical dissociations, where larger posterior and smaller anterior hippocampal volumes predict navigational proficiency in taxi drivers (Maguire et al., 2000; Woollett & Maguire, 2011) and superior recollection memory (Poppenk & Moscovitch, 2011). We extend this literature by demonstrating that representations of personal goals are also dissociated along the axis, in this context based upon their temporal distance from the present.

Examining mental constructions over time, be it past memories or future simulations, is methodologically challenging. Changes in temporal distance are inherently accompanied by changes in the properties of the representations. Construal level theory posits that humans form more abstract construals of distal entities than they do of proximal entities, where construal level refers to the degree of concreteness or abstractness of the mental representation (Liberman & Trope, 2008, 2014; Trope & Liberman, 2003, 2010). High level construals are abstract, coherent, and schematic representations and are associated with distal objects, actions, or contexts. Low level construals contain concrete, idiosyncratic, and incidental information and are assigned to proximal entities (Trope & Liberman, 2003, 2010). As a result, it becomes difficult to disentangle neural and behavioral effects arising from increases in temporal distance from those induced by changes in the nature of the constructions. We experimentally removed differences in construal level by holding the content, realness, and level of detail associated with our goals constant. We selectively varied the temporal distance of the goals and our results demonstrate that the temporal distance alone is sufficient to elicit hippocampal long axis dissociations. It is likely that when we organize our goals in the past or future, having the goals in mind in an abstract way is sufficient, whereas when we are in the present, we need more details of the goal to be able to achieve that goal.

Cortical and subcortical regions outside the hippocampus are also heavily involved in goal cognition. For instance, the ventromedial prefrontal cortex, including the medial orbitofrontal cortex and ventral medial cortex, represents the relative values of different goals (Gläscher et al., 2009; O’Doherty, 2011). The ventral striatum processes situations that are goal relevant for individuals (Montagrin et al., 2018) and computes prediction errors, i.e., deviations from reward expectations, which service goal directed behaviors by supplying updated value estimates of the world (Guo et al., 2016; Hare et al., 2008). These bodies of work have examined how the brain encodes the values of different goals, ranks possible outcomes, and selects actions that will lead to goal attainment. We distinguish ourselves from these studies by examining how the brain represents the timestamp associated with the same goals as the goals wander though the past, present, and future.

We found that current goals were processed more quickly than temporally removed goals. We theorize that this dissociation reflects the preferential status given to current needs over those that are removed, and the additional time required to mentally travel in time to place the past and future goals on a timeline. This observation is consistent with reaction time distance effects observed in temporal judgment tasks, where participants take longer to make judgments from a past or future timepoint versus a present timepoint (Arzy et al., 2009; Gauthier & van Wassenhove, 2016). Further, current goals activated more posteriorly and temporally removed goals more anteriorly in the left hippocampus.

We speculate that the timestamp of past and future goals may be localized in the anterior hippocampus in order to be integrated into schemas characterizing the time periods when the goals were already accomplished or must be accomplished (Trope and Liberman, 2003). Projecting into the future does not require to access the details of the goal, but rather an abstract representation (Conway et al; 2019; Conway 2005; 2009). Similarly, for a past goal, there is no need for specific details, its general representation is sufficient to remember the status of the goal (e.g., achieved). Furthermore, the findings of remote (anterior) versus current (posterior) representation of goals in the hippocampus in our study are in line with animal studies showing different spatial scales along the longitudinal axis of the hippocampus, with great distances (> 10 m) represented in the anterior (ventral) part, and smaller distances (< 1 m) represented in the posterior (dorsal) part (Kjelstrup et al., 2008; Poppenk et al., 2013; Strange et al., 2014). Thus, it is possible that an analogue memory temporal structure is represented in the human brain. However, more studies are needed to confirm these hypotheses.

The anterior hippocampus has preferential access to the brain’s motivational and schema circuitry through connections with the amygdala, nucleus accumbens, insula, vmPFC, perirhinal cortex, and ventral tegmental area (Grady, 2020; Kahn et al., 2008; Krebs et al., 2011; Libby et al., 2012). The anatomical distinction observed may reflect the need for past and future goals to be placed on a personal timeline and monitored in relation to other episodic and autobiographical events in one’s life (D’Argembeau, 2020). Finally, Thorp et al. (2022) suggested that taking into account the anterior medial and lateral parts of the hippocampus might bring new insights into a possibly non-linear representation of the granularity along its axis.

We found that goals were mapped based on their temporal distance in the left hippocampus, but not in the right hippocampus. We had no a priori assumptions about left and right hippocampal lateralization, but our findings align with previous work citing lateralized hippocampal activation. The right hippocampus has been more heavily involved in allocentric spatial memory and navigation (Burgess et al., 2002; Ezzati et al., 2016; Gauthier et al., 2020; Iglói et al., 2010; Smith & Milner, 1981; Spiers et al., 2001) and the left hippocampus in egocentric associative and sequential memory (Frisk & Milner, 1990; Iglói et al., 2010; Kumaran & Maguire, 2007; Maguire, 2001; Schendan et al., 2003; Spiers et al., 2001). Participants in our task were tracking goals that needed to be sequentially accomplished and were processing the goals in relation to their present selves, thus envisioning the goals from an egocentric point of view. Our hippocampal lateralization contributes to the notion that the two hippocampi provide complementary representations of the world, rather operating via a single unified function (Igloi et al. 2010).

A limitation of this research is that participants could have memorized the stimuli and relied on mnemonics to answer the questions, rather than incorporating time into their evaluations of the goals. We thoroughly trained the participants on the mission framework to reduce this risk, and we heavily emphasized during the training that it was important for them to think about when in time they needed to accomplish each goal in relation to their temporal position in the game. If participants were solely remembering sequences, we would not expect to see behavioral and neural differences between the same goals as they changed relevancy over time. Further, naturalistic goals are often learned about more incidentally and more strongly integrated into ones autobiographical knowledge base. As discussed above, we chose our space-themed design to tightly control the experiential properties of the goals and investigate the impact of the timestamp attached to a goal alone. Additional studies are warranted to examine whether existing goals from participants’ personal pasts and futures stratify to the same degree along the hippocampal long axis.

This research demonstrates that temporal distance plays a key role in guiding representations of personal goals, and this work has important implications for psychiatric disorders, namely depression. Depressed individuals generate more conflicting goals (Emmons & King, 1988), envision more obstacles to achieving their goals and form less specific goals (Vincent et al., 2004), feel less control over goal outcomes (Dickson et al., 2011), and are more pessimistic regarding the likelihood of achieving their goals (Dickson et al., 2016). It is unclear whether there are inherent differences in how depressed individuals represent the temporal distance to different goals. It is possible that distance miss-representations contribute to differences in the perceived likelihood of success and specificity of future goals. If so, re-adjusting individuals’ representations of time may offer therapeutic potential.

In conclusion, we demonstrate that the brain separates goals that are relevant for ones’ current needs from those that are removed in time. Behaviorally, this is reflected in the reaction time differences to process current and temporally removed past and future goals. Neurally, these sets of goals are dissociable along the posterior and anterior part of the hippocampal axis. We demonstrate that temporally removed goals are represented more anteriorly in the left hippocampus and current goals more posteriorly. Altogether, we provide compelling evidence for a novel application of the hippocampus’s long-axis system, where the timestamp attached to a goal dissociates its anterior and posterior organization in the hippocampus. These findings also provide new insights into how time is represented in the human brain.

## Materials and Methods

### Participants

Thirty-four healthy participants completed the experiment from New York, NY, USA. Two participants were excluded for exaggerated head motion during the scan (> 2.5mm voxel size) and one participant was excluded for technical difficulties during acquisition. Thirty-one medically and psychiatrically healthy adults were included in the final analyses (Age; M=27.0, SD=4.5, Range=19-36, n=15 females). All participants provided written informed consent and were financially compensated for their participation. The Institutional Review Board at the Icahn School of Medicine at Mount Sinai approved this experiment. Compliance with all relevant ethical regulations was ensured throughout the study and methods were carried out in accordance with relevant guidelines and regulations.

### Materials

Participants filled out 2 in-house developed screening questionnaires and a standard PDSQ (Psychiatric Diagnostic Screening Questionnaire). The first questionnaire screened for pregnancy and any neurological, psychiatric, or substance use disorders. The second questionnaire screened for magnetic resonance imaging incompatibilities. Participants completed a post-task questionnaire after the scan.

### Procedures

The experiment took place over two days. Participants completed three screening questionnaires and a training paradigm on the first day (∼1.5h). They returned to the laboratory 24 hours later for their MRI scan. Participants were first familiarized with the scanning environment and experimental task in a mock scanner during this visit (∼30m). They then underwent their 7T scan, where they completed the Mars task (∼1h), and filled out a post-task questionnaire after exiting the scanner (∼15m). See below for a comprehensive description of the training, mock scanner, and experimental paradigms.

#### Training paradigm

On the first day of the experiment, participants learned which goals they needed to accomplish in their 1^st^, 2^nd^, 3^rd^, and 4^th^ years on Mars, which goals they always needed to accomplish, and which goals they never needed to accomplish. Goals fell under the categories of (1) maintaining a part of the space shuttle, (2) maintaining a part of their spacesuit, (3) eating a certain food, (4) completing a certain exercise, and (5) enjoying a recreational activity. There were 5 goals associated with each of the 6 timeframes (year 1, year 2, year 3, year 4, always, never) totaling to 30 stimuli. Participants learned the goals using a view and quiz procedure. During the viewing phase, the goals relevant for year 1 were shown for 3000ms each, followed by a 1000ms fixation cross. During the quizzing phase, participants were randomly shown the 5 goals for year 1, as well as several foil images (i.e., goals not belonging to year 1). If the goal presented belonged to year 1, participants had to press “1” and if the goal was a foil they had to press “0” on the keyboard. Participants had 6000ms to respond and received feedback regarding whether they were correct for 1500ms, followed by a 1000ms fixation cross. If participants answered incorrectly or did not answer in time the quiz repeatedly restarted until they answered all questions correctly. Participants then advanced to viewing the 5 goals for year 2. During the year 2 quiz, participants were shown goals from year 1, year 2, and several foil images. They had to indicate “1” if the goal presented belonged to year 1, “2” if the goal presented belonged to year “2”, or “0” if the image presented was a foil. This pattern continued such that participants were cumulatively quizzed on which years each of the goals belonged to.

Participants learned the goals in the following order: year 1, year 2, year 3, year 4, always, never in the first round of the training paradigm. They then completed the training in the opposite order to ensure that all goals were quizzed equally. The first 4 quizzes in each round contained between 2-4 foil images, the last 2 quizzes per round did not contain foils. Goals were randomly presented during both the viewing and quizzing phases. The training was programmed with E-prime 2.0 (https://pstnet.com) and stickers labeled “1”, “2”, “3”, “4”, “A”, “N”, and “0” covered keys S-K on the keyboard. Participants took approximately 45-90 minutes to complete this exercise. See **Supplementary Table 6** for the instructions read and presented on the screen to the participants. The experimenter remained in the room to clarify any questions.

#### Mock scanner training

Participants were brought to a mock scanning room to be familiarized with the scanning environment and experimental task prior to their scan on the second day of the experiment. The experimenter explained to the participants that they would be embarking on a mission to Mars, and that once on Mars, the mission will last four years where they would be presented with the goals that they learned the day prior and will be asked when they need to accomplish each goal. Options included currently, in the near or distant future, in the near or distant past, always, or never. Near future was defined to the participants as 1 year in the future and distant future defined as 2 or 3 years in the future. Likewise, near past was defined to the participants as 1 year in the past and distant past defined as 2 or 3 years in the past. The experimenter emphasized that in order to succeed on the mission participants needed to think about each goal, when they had to complete it, and what Mars year they were currently in as they advanced through the mission. See **Supplementary Table 6** for the full instructions delivered to the participants during this session.

Participants were then put in the mock scanner to complete an abbreviated version of the experimental paradigm. This version navigated participants through their four years on Mars, showing them 6 sample trials per year. The stimuli presented in this version were irrelevant (the foil images from the day prior). Participants were informed that they should not focus on the content of the trial, but rather use this exercise to familiarize themselves with the graphics of the task, transition slides, and button pressing hand pads.

#### Experimental paradigm

Participants then went on their mission to Mars in the 7T scanner. The task began with a screen indicating that the participants have arrived on Mars and are starting their first Mars year. Then, they were presented with a goal and asked when they need to complete it. Using the buttons on the bottoms of the screen they indicated if they needed to complete the goal in the current year (CY), near future (NF), distant future (DF), whether they always needed to complete it (AN), or whether they never needed to complete it (NN). Responses were followed by a 0.8s or 2.5s (randomly assigned) fixation cross before the onset of the next trial. If participants were shown the space helmet, a goal that had to be completed in their 3^rd^ year in version 1 of the task, they would select the DF button to indicate that they need to complete this goal in the distant future. Participants evaluated the temporal relevance of each of the 30 goals before seeing a screen that indicated that they had succeeded in completing their 1^st^ year.

They then advanced to their 2^nd^ year on Mars and a new button, NP, appeared to indicate goals that had been accomplished in the near past. Participants re-evaluated the temporal relevance of the 30 goals, i.e., they were shown the same 30 stimuli and had to reframe their thinking now that they had moved forward in time. Continuing with the example above, participants were shown the space helmet again, which still needed to be accomplished in their 3^rd^ year. However, now that they were in their 2^nd^ year on Mars, this goal must be completed in the near future, even though last year it was something that did not need to be completed until the distant future.

When participants transitioned to their 3^rd^ year on Mars, another button, DP, appeared to indicate goals that had been completed in the distant past. The DF button was removed as there was only one year left in the mission. Participants re-evaluated the 30 goals such that the space helmet would now be a goal that must be accomplished in the current year, CY. When participants advanced to the 4^th^ Mars year, the NF button was removed as they were in their final year and there was no longer a future component. Now, the space helmet became a goal that was accomplished the previous year, so participants would select NP to indicate that they completed this goal in the near past. See **Supplementary Figure 6** for an illustration of how the game buttons changed throughout the mission.

The key aspect of this design was that the same goals were presented each year, but their temporal distance changed as the participants advanced through the game. Goals that needed to be accomplished in the 1^st^ year followed a temporal trajectory that moved from a current goal in the 1^st^ year, to a near past goal in the 2^nd^ year, a distant past goal in the 3^rd^ year, and an even more distant past goal in the 4^th^ year. Goals that needed to be accomplished in the 2^nd^ year moved from near future goals, to current goals, near past goals, and distant past goals in the 4^th^ Mars year. Goals that needed to be accomplished in the 3^rd^ year transitioned from distant future goals, to near future goals, current goals, and near past goals in the 4^th^ Mars year. And lastly, goals that needed to be completed in the 4^th^ year moved from distant future goals to distant future goals, near future goals, and current goals in the final Mars year. **Supplementary Table 7** presents a visual depiction of ^t^he temporal trajectories.

The mission contained 120 trials in total, which were divided into instances where participants were making an evaluation of a goal that needed to be completed in the distant future (15 trials), near future (15 trials), near past (15 trials), distant past (15 trials), current year (20 trials), always needed (20 trials), and never needed (20 trials). These trials were distributed across the four Mars years and a visual illustration of this distribution is provided in **Supplementary Table 1**. Participants used their index through ring fingers to press the buttons on two hand pads. The buttons rotated on the screen after every trial to prevent the reaction times for a condition from being influenced by the ease of using one finger over another. The task was self-paced, and participants did not receive feedback on their selections. Between each year, participants were given the opportunity to pause and press a button when they were ready to advance. Participants took on average 17 minutes to complete the task (M=17.79 min, SEM=0.72, Mdn=17.45 min, Range =13.08 – 33.76 min). The task was programmed in Unity (https://unity.com/)

#### Stimuli selection

Goal stimuli were selected using an independent online survey. Individuals were asked to rate the valence, arousal, and familiarity of the goals using a sliding scale from 0 [low] – 9 [high] (n=16 for the space shuttle, food, exercise, recreational activity goals, n=9 for the spacesuit goals). We visually examined that the goal ratings were neutral, i.e., no goals were viewed as extremely high or low in valence, arousal, or familiarity. Goals from each category were randomly assigned to sets, such that each set of stimuli had a space shuttle, space suit, food, exercise, and recreational activity goal. Each set of goals stayed as a unit for this point forward, but the timeframe that the goal set was assigned to (year 1, year 2, year, 3, year 4, always, never) varied by task version. We verified that there were no differences in valence, arousal, and familiarity across each set of goals (**Supplementary Figure 7**, Kruskal-Wallis rank sum test; valence: X^2^(5) =3.02, *p*=0.70, arousal: X^2^(5) =6.11, *p*=0.30, familiarity: X^2^(5) =2.05, P=0.84). The surveys were administered via SurveyMonkey (https://www.surveymonkey.com).

#### Task versions

We created 6 versions of the task such that each set of goals was associated with every timeframe (to be accomplished in year 1, year 2, year, 3, year 4, always, never). Each set of 5 goals stayed as a unit. However, a set needed to be accomplished in year 1 for participants completing version 1 of the task, in year 2 for participants completing version 2 of the task and never for participants completing version 6 of the task. See **Supplementary Figure 8** for an illustration of the stimuli and version system and **Supplementary Table 8** for a distribution of participants by version.

#### Additional choice trials

Randomly throughout the mission, participants were presented with two goals on the screen and instructed to choose the goal they felt would most benefit their survival at the present moment. The choice was between a goal relevant for the current year and a goal relevant for another timeframe, and participants indicated their choice by clicking either the “Left” or “Right” thumb buttons. They were presented with 100 choices in total (25 per year), with a current item being compared to a distant (30 trials) future (15 trials), near future (15 trials), near past (15 trials), distant past (15 trials), always needed (20 trials), and never needed (20 trials) goal (see **Supplementary Table 5** for results).

#### Post-task questionnaire

Participants filled out a questionnaire immediately after exiting the scanner. This questionnaire asked participants whether they were hungry during the scan, motivated to complete the task, whether motivation lessened at any point during the task, whether they employed a strategy to remember the goals, whether they felt time was passing during the mission, and lastly, a free recall task (to write down as many goals as they could remember). An analysis of this data is not included.

### Neuroimaging methods

#### fMRI acquisition

Data was acquired on a 7T Siemens Magnetom scanner (Erlangen, Germany) with a 32-channel head coil (Nova Medical, Wilmington, MA). Anatomical images were collected with a twice magnetization-prepared rapid gradient echo (MP2RAGE) sequence for improved T1-weighted contrast and spatial resolution (Marques et al., 2010) [0.7mm isotropic resolution, 224 slices, TR=6000ms, TE=5.14ms, field of view=320x320, bandwidth=130 Hz/Px]. Functional data was acquired in a single run using a multi-echo multi-band echo-planar imaging (EPI) pulse sequence [2.5mm isotropic resolution, 50 slices, TR=1850ms, Tes= [8.5, 23.17, 37.84, 52.51 ms], MB=2, iPAT acceleration factor=3, flip=70, field of view=640x640, pixel bandwidth=1786]. This acquisition approach situated us in a prime position to detect small biological effects. Ultra-high field 7T fMRI provides 2x the signal-to-noise ratio as traditional 3T fMRI (Kundu et al., 2017), and multi-echo protocols maximize BOLD contrast throughout the brain and offer 5-fold increases in statistical power over singe-echo acquisitions (Kundu et al., 2012, 2017).

#### fMRI data preprocessing

Functional files were denoised for physiological and motion artifacts using multi-echo independent components analysis (ME-ICA, https://bitbucket.org/prantikk/me-ica). A detailed description of this pipeline can be found here (Kundu et al., 2012, 2017). In short, ME-ICA is based on the observation that BOLD signals have linearly TE-dependent percent signal changes, as a result of T2* decay, while artifacts do not. ME-ICA decomposes multi-echo functional data into independent components and computes the TE dependence of the BOLD signal for each component. It categorizes components as either BOLD or non-BOLD and removes the non-BOLD components, leaving the data robustly denoised of motion, physiological, and scanner artifacts. As shown in Morris et al. (2019), the use of the ME-ICA pipeline further enhances the signal power gains obtained when moving from 3T to ultra-high field 7T fMRI (Morris et al., 2019).

The ME-ICA pipeline outputted a denoised timeseries with T1 equilibration correction (including thermal noise) in the subject’s native space and the remaining preprocessing steps were performed using the Functional MRI of the Brain Software Library (FSL) version 5.0.10 (Jenkinson et al., 2012). Anatomical images were first skull-stripped using robust brain centre estimation in BET. Functional images were registered to the subject-specific high resolution T1-weighted structural images using boundary-based registration (Greve & Fischl, 2009) and to a standard brain image (MNI 152 T1 2.5mm^3^) using a 12 DOF linear registration in FSL’s fMRI Expert Analysis Tool, FEAT (Jenkinson et al., 2002). Since the task was self-paced and the scans manually stopped, registered functional files were trimmed to include only 7 volumes following each subject’s completion of the task. Images were spatially smoothed at 5mm FWHM (double the voxel size to ensure GRF theory validity and this is the most appropriate smoothing for smaller structures such as the hippocampus (Poldrack et al., 2011; Mickl et al., 2008).

#### Regions of interest

We defined individual right and left hippocampal masks using FreeSurfer v7.2, we segmented the hippocampal subfields and nuclei of the amygdala (Iglesias et al., 2015). We first used the ‘recon-all’ and ‘autorecon1’ functions to get the t1 whole brain scan for each participants. Then we used the hippocampus segmentation tool (‘segmentHA_t1.sh’), we reoriented the masks using ‘fslreorient2std’, we merged the different parts of the hippocampus (i.e., CA1, CA2, etc.) using ‘fslmaths’. Masks were registered applying the transformation estimated between the t1 and the 2.5mm^3^ brain template using FSL (‘flirt’ and ‘applyxfm’). Finally, the group average ROI for hippocampus subportions was obtained by binarizing and averaging individually drawn ROIs for all subjects, and selecting only voxels common to at least 80% of the participants. We created a whole brain gray matter mask by resampling SPM12’s gray matter tissue probability map to the 2.5mm^3^ space and thresholding with a probability threshold of 20%. **Supplementary Figure 9** visualizes all masks.

### Analyses

#### Behavioral analyses

Participants’ reaction time data was analyzed using a series of linear mixed models, implemented using the lme4 and lmerTest packages in R v. 3.6.0. The data was first processed to remove trials where the participant got the question incorrect, and trials considered outliers (reaction times +/- 3 SD’s of a participant’s mean). This left 94.5% of all trials across the 31 participants. Reaction times were log transformed prior to modeling. The first model examined the relationship between participants’ log transformed reaction times as the dependent variable and temporal condition as the independent variable. Temporal condition was modeled as a fixed effect with five levels: distant future, near future, near past, distant past, and current trials. We removed always and never trials from the data set for this model because they inherently differed in that they consisted of the same stimuli each game year, while the stimuli in the other categories updated as time progressed.

We validated that the results held in a second model that included the always and never trials. The second model examined the relationship between participants’ log transformed reaction times as the dependent variable and trial type as the independent variable. Trial type was modeled as a fixed effect with two levels. Distant future, near future, near past, and distant past trials were coded as those involving a temporal component, while current, always, and never trials were coded as those not.

Participant ID was included as a random effect to account for the lack of independence between trials within a participant, and game year was included as a fixed effect and as a random effect for objective time (game year) to account for trends in reaction time across the years as the mission proceeded within and between participants. This game year regressor had 4 levels (year 1, year 2, year 3, year 4) and was in place to absorb variance such that any reaction time differences that remained between the temporal conditions were those above and beyond what could be explained by objective time point in the game.

Statistical significance was evaluated using Type II Sums of Squares. Model-based estimated marginal means, standard errors, confidence limits, and *post hoc* tests were computed via the emmeans package in R. *Post hoc* tests compared two estimated marginal means and outputted the corresponding estimate, standard error, test statistic, and p value. Degrees of freedom were approximated using the Satterthwaite method and *post hoc* tests were corrected for multiple comparisons using a Tukey adjustment. See **Table 1** for the model formulas.

#### General linear modeling analyses

Univariate analyses consisted of general linear models implemented in FSL’s FEAT (Jenkinson et al., 2012).

#### Primary analysis

For the first-level analyses, models included four separate regressors, one for each temporal condition, including only correct responses: remote, current, always, and never.

#### Secondary analysis

For the first-level analyses, models included seven separate regressors, one for each temporal condition, including only correct responses: distant future, near future, near past, distant past, current, always, and never.

#### Primary and secondary analyses

Regressors for each game year (year 1, year 2, year 3, year 4) were also included in the model. These contained all trials in the year specified, i.e. the task, additional choice, incorrect responses, and reaction time outlier trials were in place to absorb variance explained by position in the task. The duration of each trial was set as the participant’s reaction time on that trial due to the self-paced nature of the experiment. All trials were modeled with boxcars and convolved with FEAT’s canonical hemodynamic response function. Temporal derivatives of the aforementioned regressors and six motion regressors were also included in the model. See **Supplementary Table 9 (primary analysis) and Supplementary Table 10 (secondary analysis)** for all regressors included in the analysis. A high pass filter with a cutoff of 128s was applied to remove low frequency signal drifts.

Two contrasts (primary analysis) compared the temporally removed condition to the current condition (remote > current, and current > remote). Eight contrasts (secondary analysis) compared the temporally removed conditions to the current condition (distant future > current, near future > current, near past > current, distant past > current, and the reverse). First level activation maps were brought to a second level mixed effects analysis, implemented in FLAME 1+2 (FSL’s Local Analysis of Mixed Effects), where one sample t-tests were used to determine the group mean for each contrast. Statistical maps were masked as described above and results corrected for multiple comparisons using Gaussian random field theory (Worsley, 2001). Hippocampal and whole brain GLMs were corrected for multiple comparisons using GRF-theory-based maximum height thresholding with a family wise error (FWE) corrected significance threshold of *p*=0.025. Activation maps were visualized using MRIcroGL’s suite of visualization tools (Rorden & Brett, 2000) (https://www.nitrc.org/projects/mricrogl).

## Acknowledgements

The authors thank Dominik Moser for his help with the pilot studies, Matthew Schafer and Ofer Perl for their helpful discussions, Michaël Dayan, Ben Meulman for their help with MRI and behaviour statistics, and Yael Jacobs for her assistance implementing the fMRI preprocessing pipeline. We would also like to thank Thomas Grand for illustrating the stimuli and Adrienne Young for assisting with the Unity programming. D.S. is supported by the National Institute of Health, USA (R01MH122611, R01MH123069); A.M. is supported by a Swiss National Science Foundation Grant (P1GEP1-148618); and D.E.C. is supported by a NSF graduate research fellowship.

## Author contributions

A.M. developed the research question; A.M., D.E.C., and D.S. designed the research; L.L programmed the task; D.E.C. collected the data; A.M., and D.E.C. analyzed the data with the input from M.G.P, M.G.B, and D.S. A.M., and D.E.C. wrote the manuscript with input from all authors.

## Data availability

The dataset generated and analyzed during the current study is available in the Open Science Framework repository [https://osf.io/p5zwx/].

## SUPPLEMENTARY MATERIAL

**Supplementary Figure 1.**
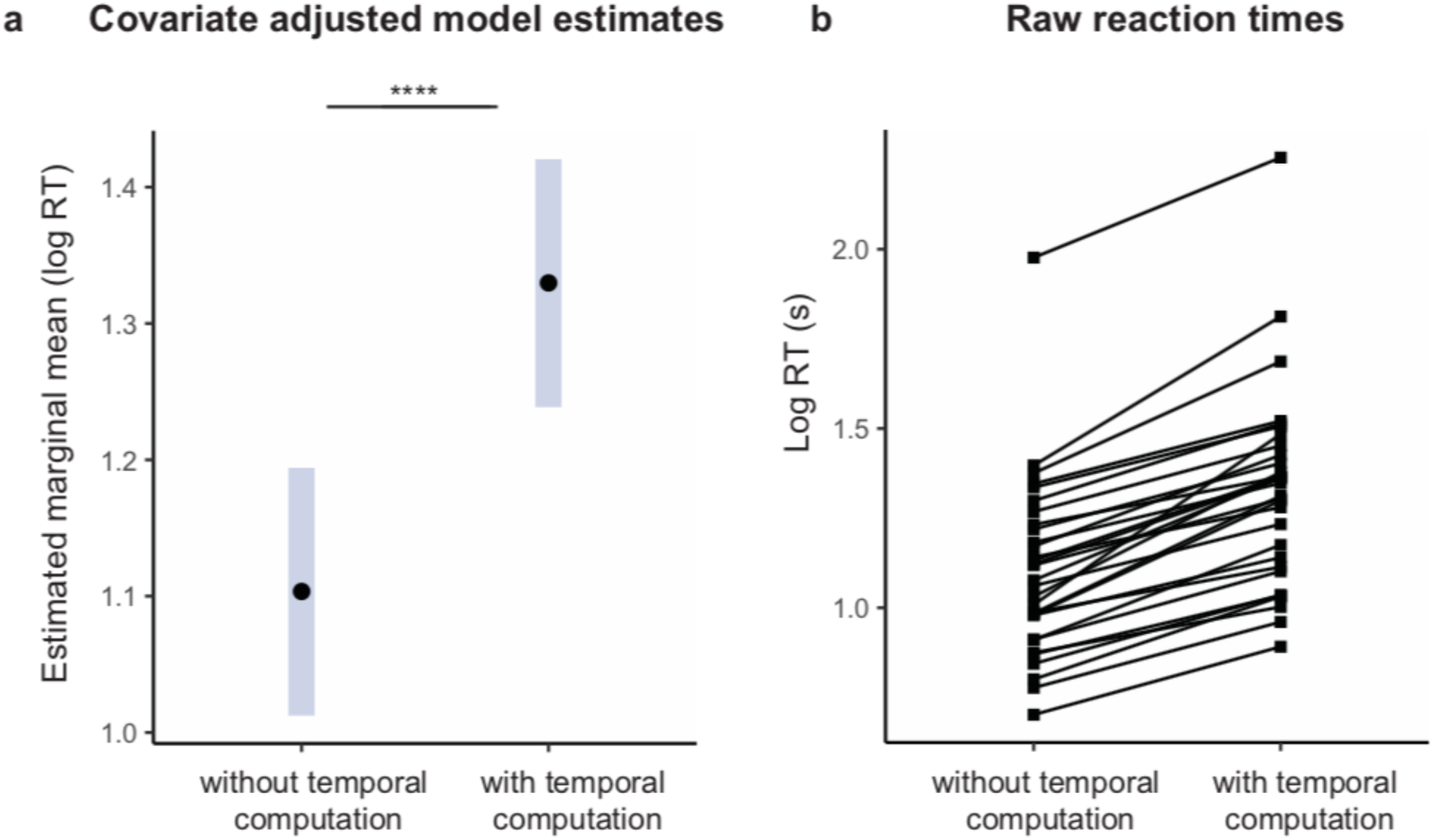
Participants took longer to process goals that involved a temporal computation. We explored the relationship between participants’ log transformed reaction times and trial type using a linear mixed model. We compared trials with a temporal computation (distant future, near future, near past, distant past) to those without a temporal computation (current, always, never). Log transformed reaction time was significantly associated with trial type (Type II ANOVA; *F*(1, 3422.7) = 455.464, *p*<0.0001). **(a)** A *post hoc* examination of the difference between the two estimated marginal means revealed that temporally removed goals took significantly longer to process (0.226 ± 0.0106, Z = 21.342, *p*<0.0001). Error bars = 95% CI. **(b)** This effect was robustly present at the individual subject level when examining participants’ average log reaction times. \*\*\*\**p*<0.0001.

**Supplementary Figure 2.**
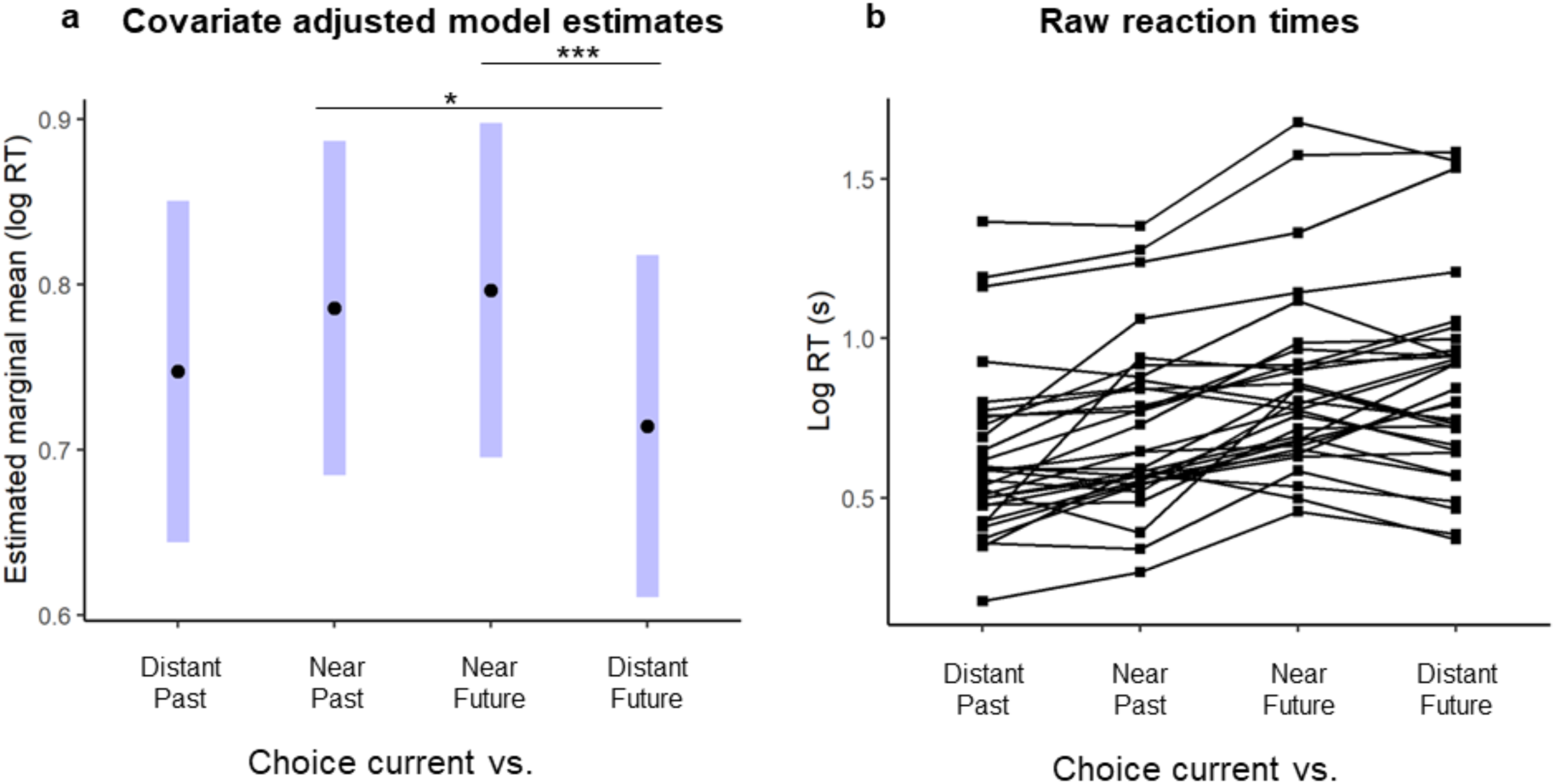
Additional choice trials (Game2): Participants took longer to choose between current and near future or near past goals than current and distant future goals. (a) *Post hoc* comparisons of the estimated marginal means revealed that when participants had to choose between current and distant future goals they processed faster than when participants had to choose between current and near future or near past. Error bars = 95% confidence Interval (CI). (b) Individual participants’ average log reaction time for each of the temporal conditions against current. *** *p*=0.001, \**p*<0.05.

**Supplementary Figure 3.**
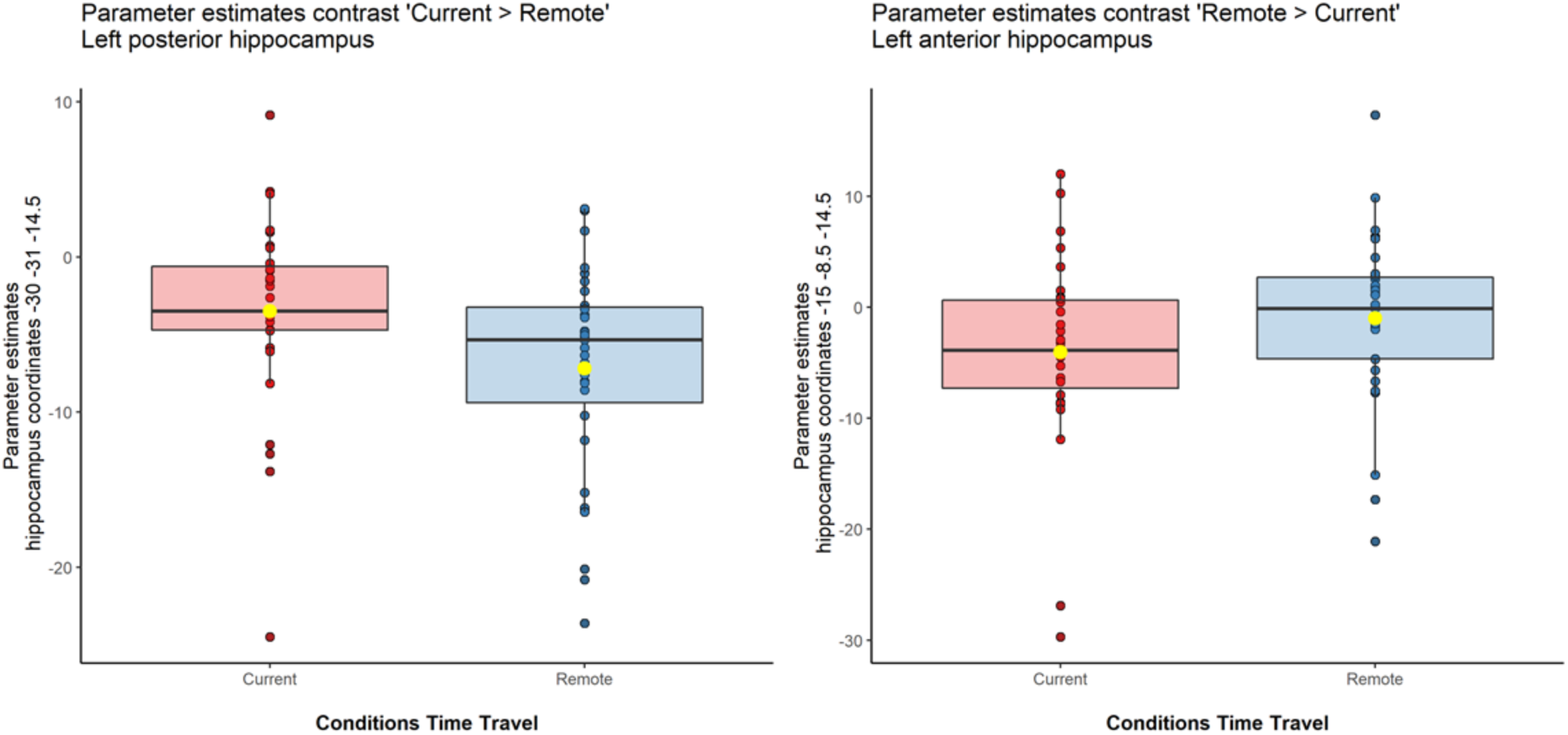
Parameter estimates of left hippocampus separately for two coordinates. On the left panel, posterior hippocampus coordinates extracted from first level for the condition “Current” and “Remote” from the peak voxel [-30 -31 -14.5], (current condition: *Mean* = -3.51, *SEM* = 1.11; remote condition: *Mean* = -7.18, *SEM* = 1.22). On the right panel, anterior hippocampus coordinated extracted from first level for the condition Remote and Current, from the peak voxel [-15 -8.5 -14.5], (current condition: *Mean* = -4.05, *SEM* .1.54; remote condition: *Mean* = -1.01, *SEM* = 1.38). Error bars = standard error of the mean (SEM).

**Supplementary Figure 4.**
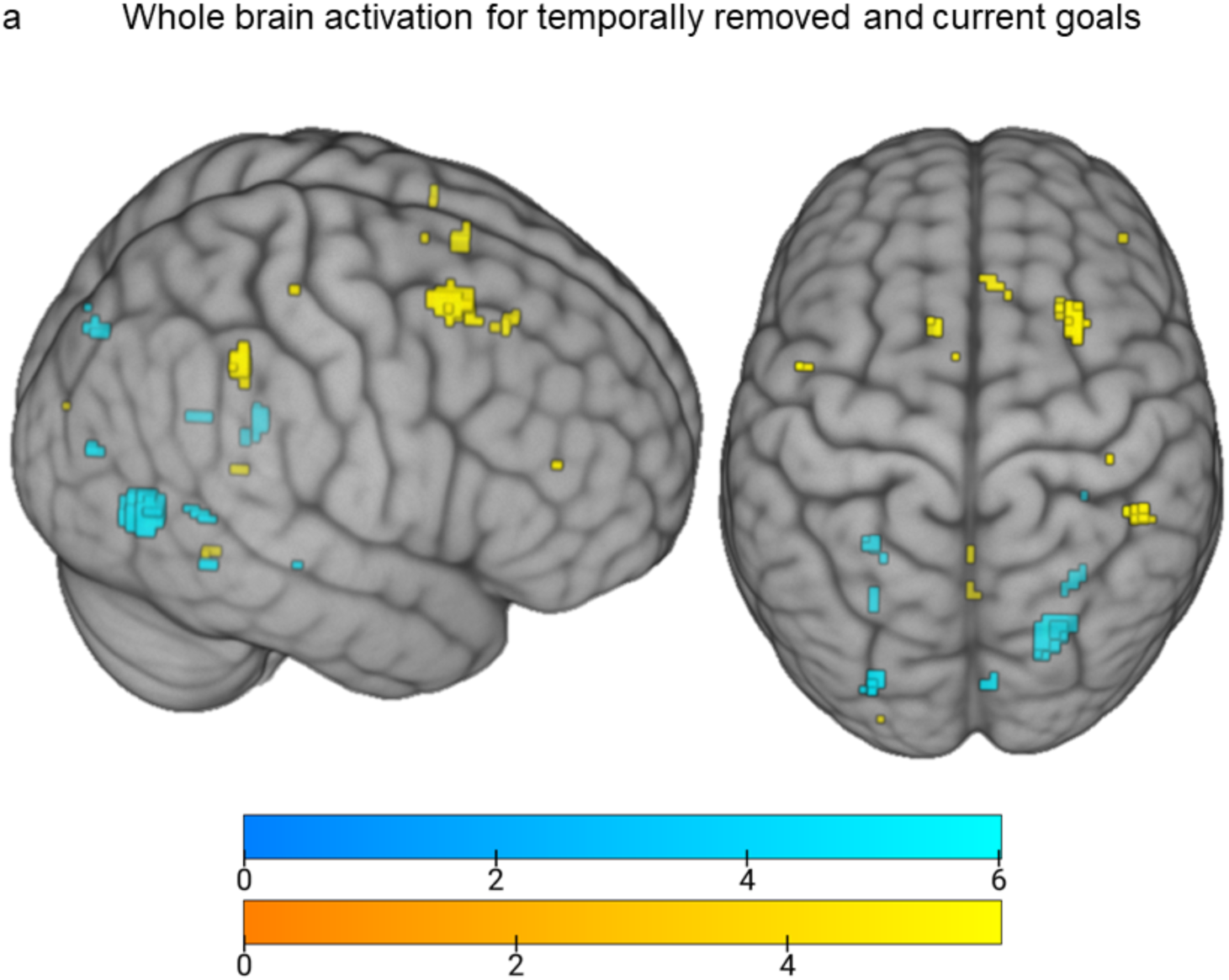
Whole brain GLM results for temporally removed and current goals. Goals that were removed in time activated more anterior regions of the brain, while current goals activated more posterior regions of the brain. **(a)** A contrast comparing the Remote (Distant + Near Future + Distant + Near Past) > Current are overlaid in yellow. A contrast comparing the Current > Remote (Distant + Near Future + Distant + Near Past) are overlaid in blue. All z-statistic images were thresholded parametrically using maximum height thresholding (FWE voxel-wise correction, *p*=0.025). Contrasts maps were overlaid and rendered onto a 3-dimensional MNI 152 brain using MRIcroGL. Color bars reflect the thresholded z-statistic scores [z=0-6].

**Supplementary Figure 5.**
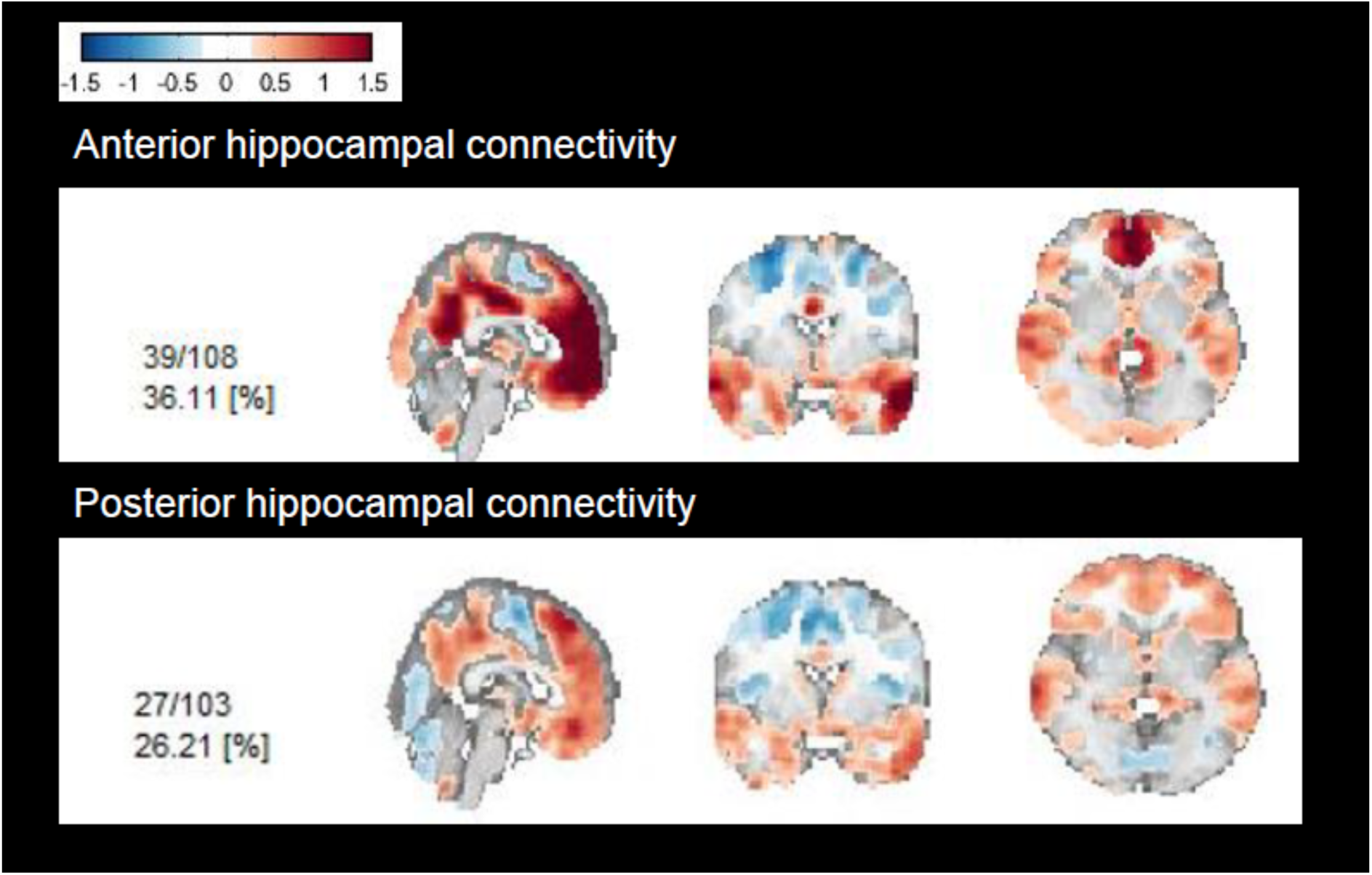
Exploratory analysis of functional connectivity. We employed seed-based co-activation patterns (CAPs; Bolton et al., 2020). with the anterior (head) and the posterior (tail+body) hippocampus as seeds in two separate analyses. We found that both anterior and posterior hippocampus co-activated with the frontal, inferior temporal, and posterior parietal cortices to different degrees (with anterior hippocampus showing overall stronger connectivity), while a clear difference was observed in the posterior occipital areas, which co-activated with the posterior but not anterior hippocampus (see figure above). These findings are intriguing given that the posterior hippocampus is structurally more connected to the posterior part of the brain (occipital and medial parietal cortex), while both anterior and posterior are structurally connected with frontal areas (Dalton et al., 2022).

**Supplementary Figure 6.**
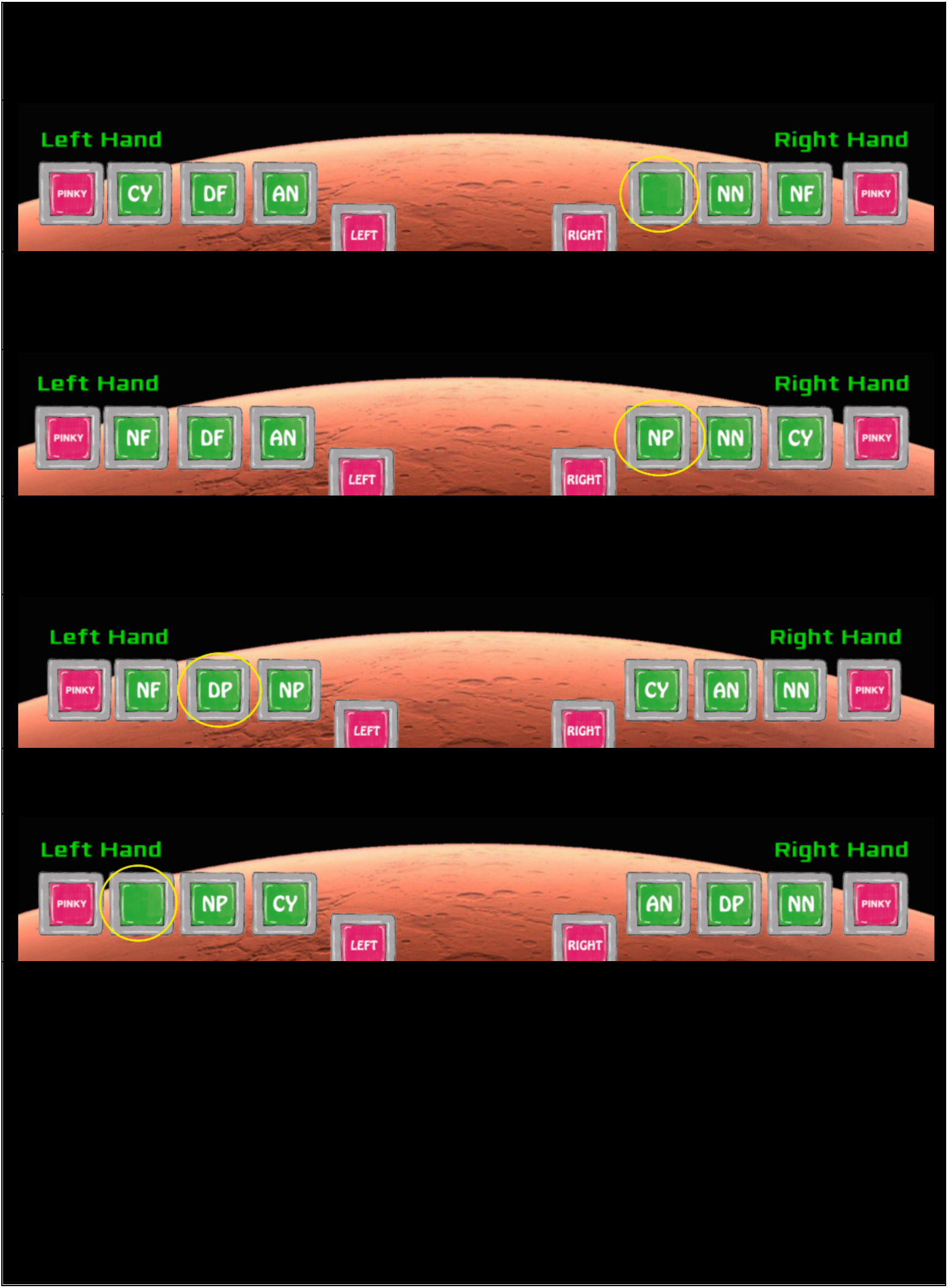
Game buttons. Participants selected their responses using hand pads attached to their right and left hands. The button options remained the same within a year and rotated on the screen after every trial to prevent the reaction times for a condition from being influenced by the ease of using one finger over another. Across years, the buttons present on the screen changed depending on whether past or future options were applicable.

**Supplementary Figure 7.**
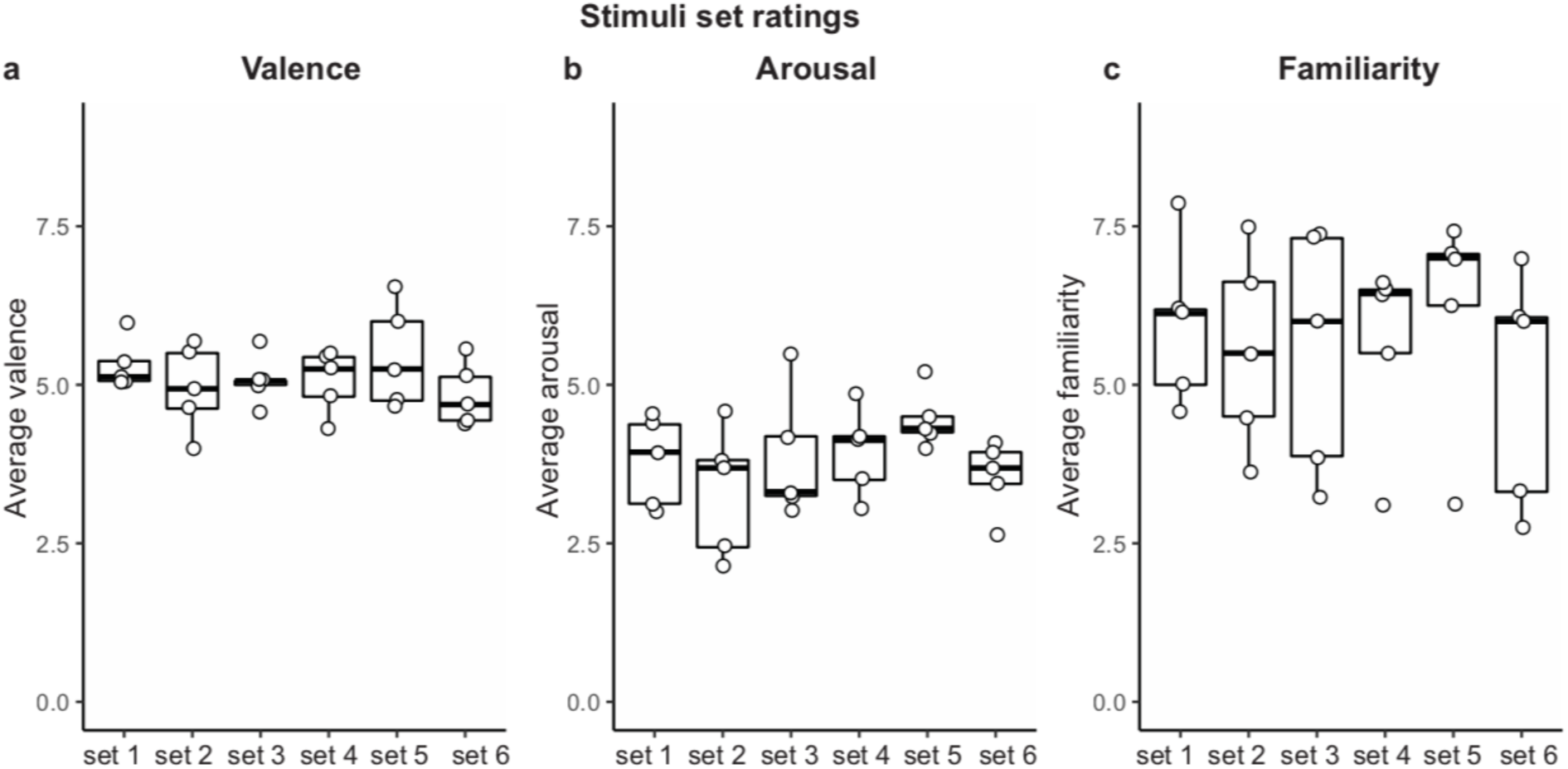
Stimuli selection. Goals were selected using an online survey in an independent sample. The goals selected belonged to 5 categories (space shuttle, space suit, food, exercise, and recreational activity) and one goal from each category was randomly assigned to a set. There were no differences in **(a)** valence (Kruskal-Wallis rank sum test; X^2^_(5)_ =3.02, *p*=0.70) **(b)** arousal (Kruskal-Wallis rank sum test; X^2^_(5)_ =6.11, *p*=0.30) or **(c)** familiarity (Kruskal-Wallis rank sum test; X^2^_(5)_ =2.05, *p*=0.84) across each set of goals. Each goal set stayed as a unit. However, the timeframe that the goal set was assigned to (year 1, year 2, year, 3, year 4, always, never) varied by task version. Box plots; center line=median, box limits=Q1 and Q3, whiskers= smallest/largest value no further than 1.5x IQR. Individual data points= average rating for each goal stimulus in the set.

**Supplementary Figure 8.**
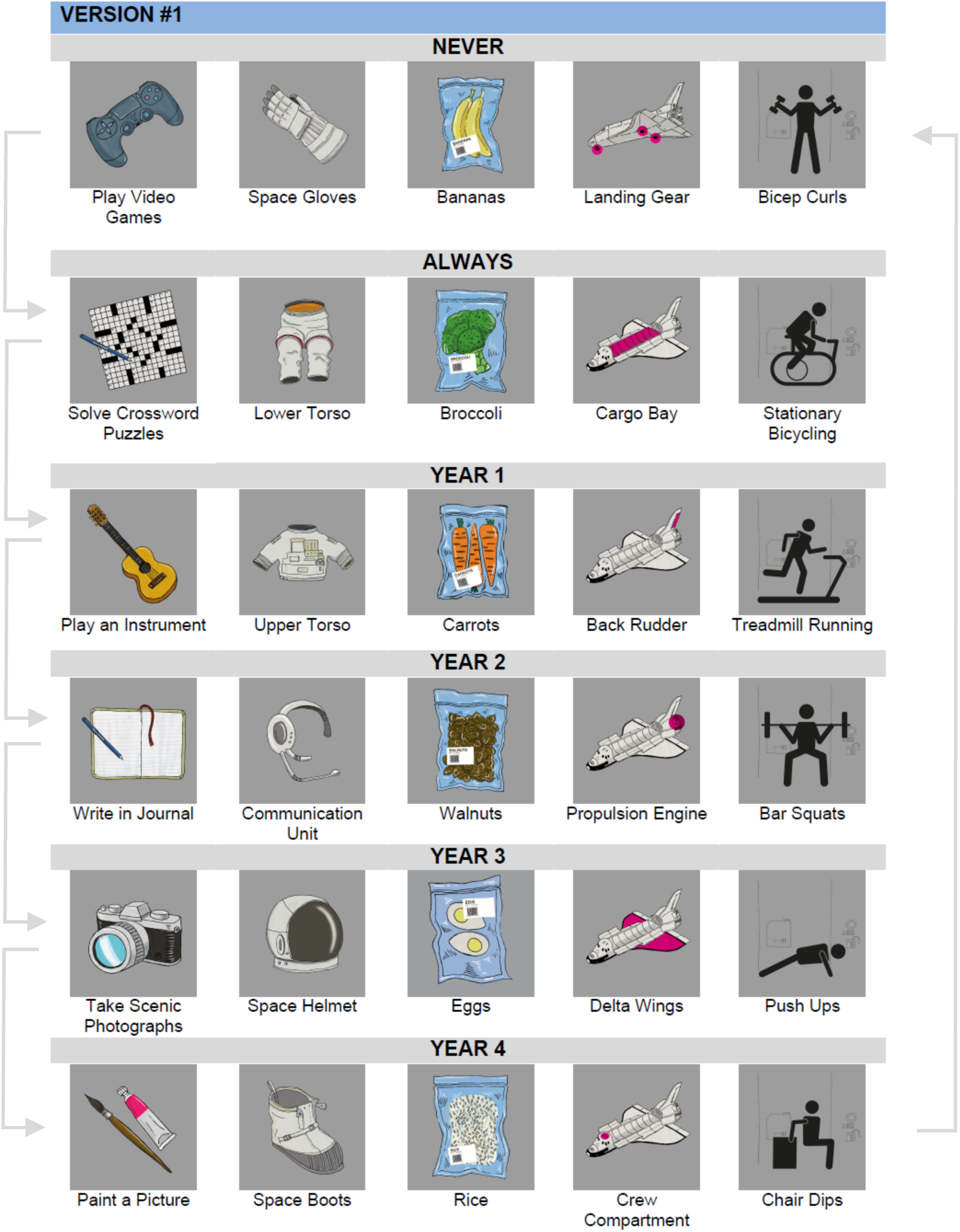
Stimuli and version visualization. Stimuli and timeframe assignments for version 1 of the task. The arrows indicate how the stimuli transitioned across the 6 versions of the task. Each set of 5 goals stayed as a unit and was assigned to every timeframe (year 1, year 2, year 3, year 4, always, never) across the different versions.

**Supplementary Figure 9.**
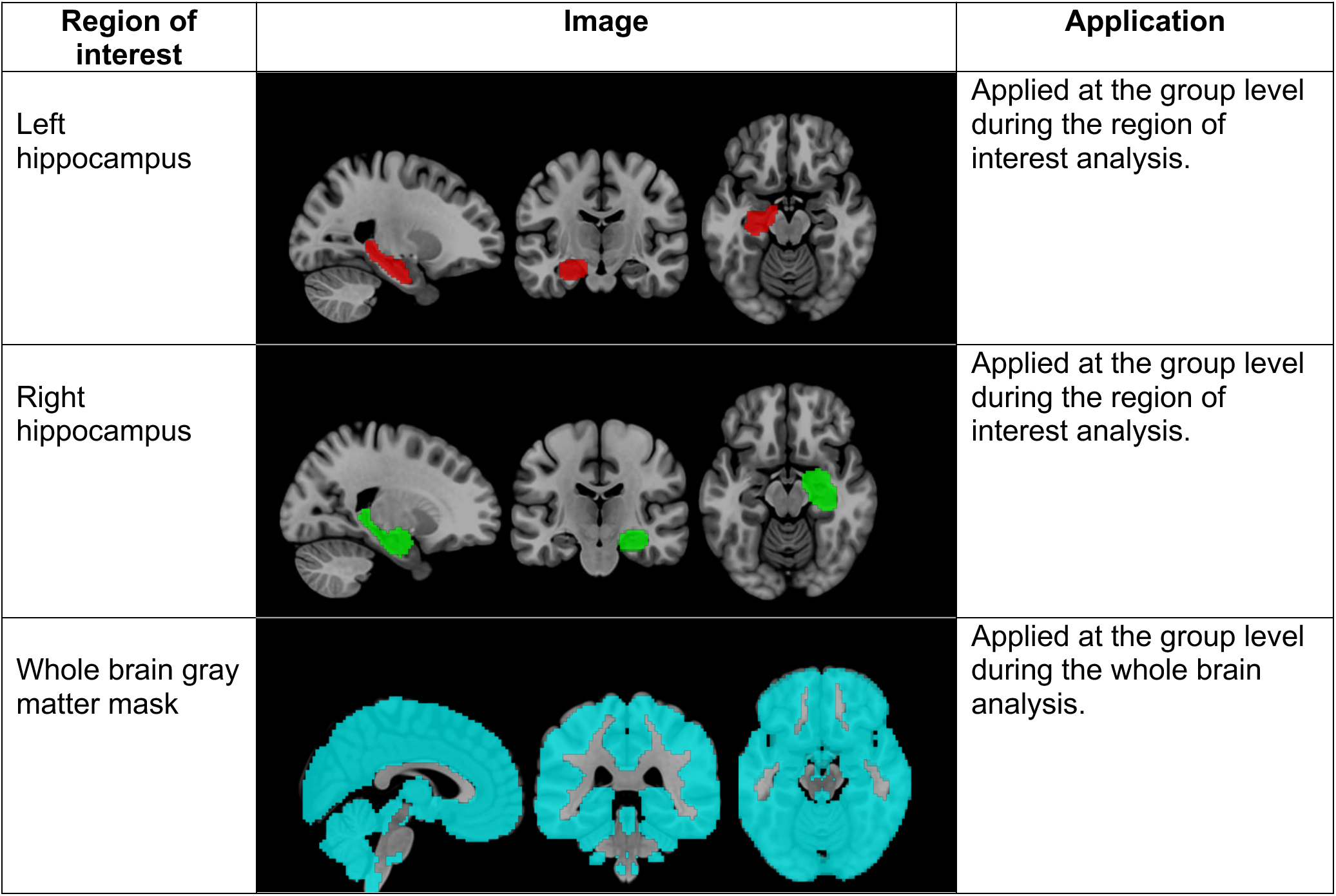
Functional MRI masks. All masks are in the 2.5mm^3^ MNI space and overlaid a standard brain image, MNI 152 T1 2mm^3^.

**Supplementary Table 1.** Distribution of game trials. Participants were presented with 120 trials during the game. The numbers in the boxes reflect the number of trials for the given temporal condition during that game year. There were 20 trials each for the current, always, and never conditions, and 15 trials each for the temporally removed conditions. Current, always, and never trials were presented across all four game years. Distant future trials were presented in the first two game years, near future trials presented in the first three, near past trials presented in the last three, and distant past trials presented in the last two game years.

|  | Mars year 1 | Mars year 2 | Mars year 3 | Mars year 4 |
| --- | --- | --- | --- | --- |
| Current | 5 | 5 | 5 | 5 |
| Always | 5 | 5 | 5 | 5 |
| Never | 5 | 5 | 5 | 5 |
| Distant future | 10 | 5 |  |  |
| Near future | 5 | 5 | 5 |  |
| Near past |  | 5 | 5 | 5 |
| Distant past |  |  | 5 | 10 |

**Supplementary Table 2.** *Post hoc* reaction time comparisons for current versus temporally removed goals. We modeled participants log transformed reaction times against temporal condition, with levels for distant future, near future, current, near past, and distant past trials. Participants processed current goals more quickly than all temporally removed past and future goals when compared individually via *post hoc* contrasts. Estimated marginal means and standard errors for the temporal conditions were as follows: distant future (1.30 ± 0.049), near future (1.34 ± 0.048), current (1.11 ± 0.048), near past (1.41 ± 0.048), distant past (1.28 ± 0.049). The estimates above represent *post hoc* comparisons of the estimated marginal means for each condition. The *p* values were corrected for multiple comparisons using the Tukey method for comparing a family of 5 estimates. Degrees of freedom were approximated using the Satterthwaite method.

| Contrast | Estimate | SE | df | t.ratio | P value | Significance |
| --- | --- | --- | --- | --- | --- | --- |
| current vs. distant future | -0.186 | 0.021 | 3419.8 | -8.81 | <0.0001 | *** |
| current vs. near future | -0.232 | 0.020 | 3417 | -11.603 | <0.0001 | *** |
| current vs. near past | -0.297 | 0.02 | 3414.7 | -14.87 | <0.0001 | *** |
| current vs. distant past | -0.166 | 0.021 | 3414.3 | -8.06 | <0.0001 | *** |

**Supplementary Table 3.** Coordinates of peak activation for each temporally removed and current goals in the left hippocampus. Each temporally removed goals (Distant Future, Near Future, Near Past and Distant Past) activated the left anterior hippocampus, while current goals activated the left posterior hippocampus (except Current > Distant Future that show no activation). This table reports the number of significant voxels in the cluster, maximum z-statistic within the cluster, and the x, y, z location of the maximum intensity voxel for each contrast. Coordinates are reported in Montreal Neurological Institute (MNI) space. All statistical maps were corrected for multiple comparisons using maximum height thresholding (FWE voxel-wise correction, *p*=0.025).

| Contrast | Voxel size | Hemisphere | Hippocampus Axis | Z-stat | Coordinates in mm (MNI) |  |  |
| --- | --- | --- | --- | --- | --- | --- | --- |
| Current > Distant Future | - | - | - | - | - | - | - |
| Current > Near Future | 3 | L | Posterior | 4.13 | -32.5 | -26 | -19.5 |
| Current > Near Past | 11 | L | Posterior | 4.04 | -30 | -31 | -14.5 |
| Current > Distant Past | 1 | L | Posterior | 3.56 | -32.5 | -26 | -19.5 |
| Distant Future > Current | 2 | L | Anterior | 3.73 | -12.5 | -13.5 | -22 |
| Near Future > Current | 1 | L | Anterior | 3.55 | -17.5 | -11 | -14.5 |
| Near Past > Current | 14 | L | Anterior | 4.6 | -15 | -8.5 | -14.5 |
| Distant Past > Current | 10 | L | Anterior | 4.28 | -12.5 | -8.5 | -14.5 |

**Supplementary Table 4.** Whole brain activation coordinates for temporally removed goals. Temporally removed (past and future) goals activated bilaterally more anterior regions of the brain, while current goals activated bilaterally more posterior regions of the brain. This table reports the number of significant voxels in the cluster, maximum z-statistic within the cluster, and the x, y, z location of the maximum intensity voxel for each contrast. Coordinates are reported in Montreal Neurological Institute (MNI) space. All statistical maps were corrected for multiple comparisons using maximum height thresholding (FWE voxel-wise correction, *p*=0.025).

| Contrast | voxel size | z-max | Coordinates in mm |  |  | Hemisphere | Lobe | Region |
| --- | --- | --- | --- | --- | --- | --- | --- | --- |
| **current > remote** | 56 | 6.02 | 22.5 | -68.5 | -9.5 | R | Occipital | Lingual Gyrus |
|  | 13 | 5.01 | -30 | -46 | -7 | L | Limbic/Occipital | Fusiform/Lingual Gyrus |
|  | 7 | 5.13 | -27.5 | -88.5 | 15.5 | L | Occipital | Occipital Gyrus |
|  | 6 | 5.25 | 30 | -53.5 | -7 | R | Occipital | Lingual Gyrus |
|  | 3 | 4.95 | -27.5 | -61 | -9.5 | L | Occipital | Lingual Gyrus |
|  | 3 | 5.02 | 7.5 | -86 | -2 | R | Occipital | Calcarine |
|  | 2 | 4.93 | 30 | -51 | -22 | R | Cerebellum | Culmen |
|  | 1 | 4.84 | 32.5 | -28.5 | -19.5 | R | Limbic | Parahippocampal Gyrus |
| **remote > current** | 33 | 5.52 | 30 | 11.5 | 58 | R | Frontal | Middle Frontal Gyrus |
|  | 18 | 5.56 | 45 | -41 | 48 | R | Parietal | Inferior Parietal Lobule |
|  | 7 | 5.17 | 7.5 | 24 | 40.5 | R | Frontal | Cingulate Gyrus |
|  | 7 | 5.32 | -10 | 11.5 | 58 | L | Frontal | Cingulate Gyrus |
|  | 3 | 4.93 | 0 | -53.5 | -34.5 | L/R | Cerebellum | Vermis |
|  | 3 | 5.57 | -50 | -1 | 50.5 | L | Frontal | Precentral Gyrus |
|  | 2 | 5.08 | 0 | -48.5 | -9.5 | L/R | Cerebellum | Vermis |
|  | 1 | 4.83 | -25 | -96 | -7 | L | Occipital | Cuneus |
|  | 1 | 4.9 | 42.5 | 41.5 | 20.5 | R | Frontal | Middle Frontal Gyrus |
|  | 1 | 4.84 | -5 | 1.5 | 60.5 | L | Frontal | Supplemental Motor Area |
|  | 1 | 4.83 | 40 | -28.5 | 68 | R | Frontal | Precentral Gyrus |

**Supplementary Table 5.** Coordinates of peak activation for never and always needed conditions against temporally removed and current goals in the left hippocampus. Always needed goals against remote goals activated the left posterior hippocampus, similarly to the activation found for current against remote. However, always needed against current goals activated the left anterior part pf the hippocampus while the never needed goals does not show any voxel active against the current condition. This table reports the number of significant voxels in the cluster, maximum z-statistic within the cluster, and the x, y, z location of the maximum intensity voxel for each contrast. Coordinates are reported in Montreal Neurological Institute (MNI) space. All statistical maps were corrected for multiple comparisons using maximum height thresholding (FWE voxel-wise correction, *p*=0.025). AN, Always Needed; NN, Never Needed.

| Contrast | Voxel size | Hemisphere | Hippocampus Axis | Z-stat | Coordinates in mm (MNI) |  |  |
| --- | --- | --- | --- | --- | --- | --- | --- |
| AN > Remote | 13 | L | Posterior | 3.95 | -35 | -23.5 | -12 |
|  | 1 | L | Anterior | 3.75 | -13.5 | -14.5 | -32.5 |
| Remote > AN | - | - | - | - | - | - | - |
| AN > Current | 19 | L | Anterior | 4.03 | -12.5 | -8.5 | -14.5 |
|  | 1 | L | Posterior | 3.57 | -32.5 | -28.5 | -14.5 |
| Current > AN | - | - | - | - | - | - | - |
| AN > NN | - | - | - | - | - | - | - |
| NN > AN | - | - | - | - | - | - | - |
| NN > Remote | - | - | - | - | - | - | - |
| Remote > NN | 175 | L | Posterior | 4.81 | -27.5 | -23.5 | -14.5 |
| NN > Current | 125 | L | Anterior | 5.01 | -17.5 | -11 | -12 |
| Current > NN | - | - | - | - | - | - | - |

**Supplementary Table 6.**
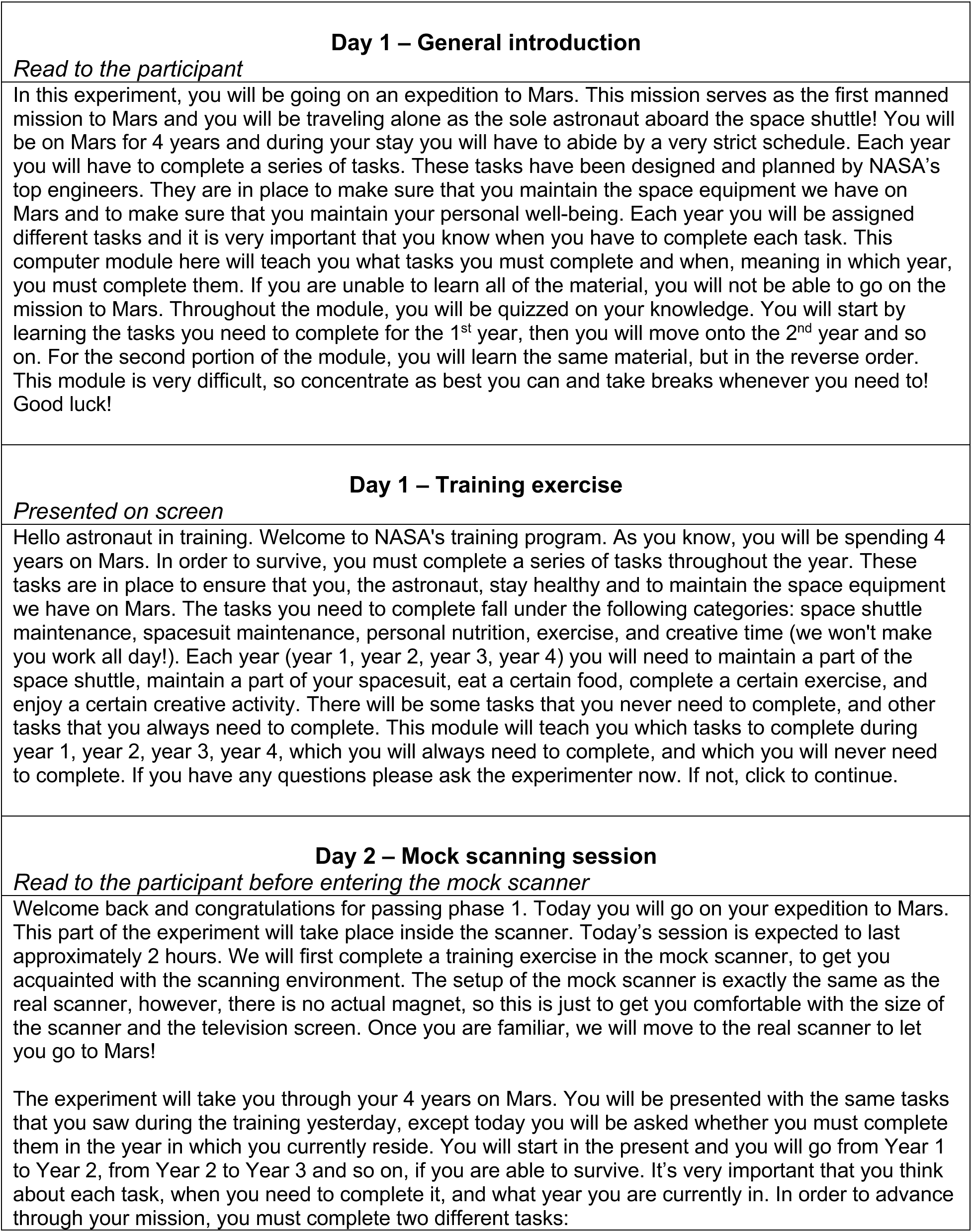
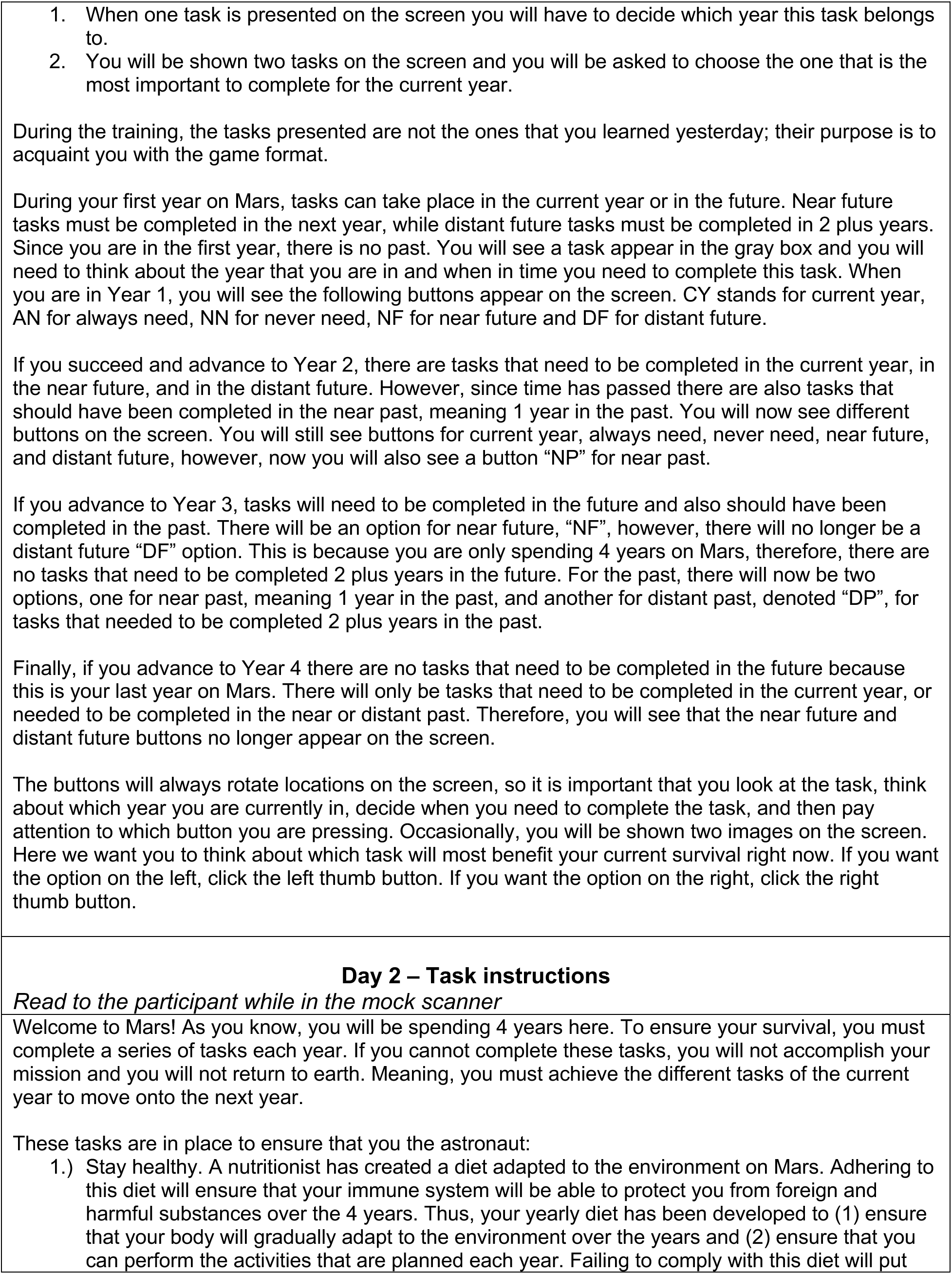
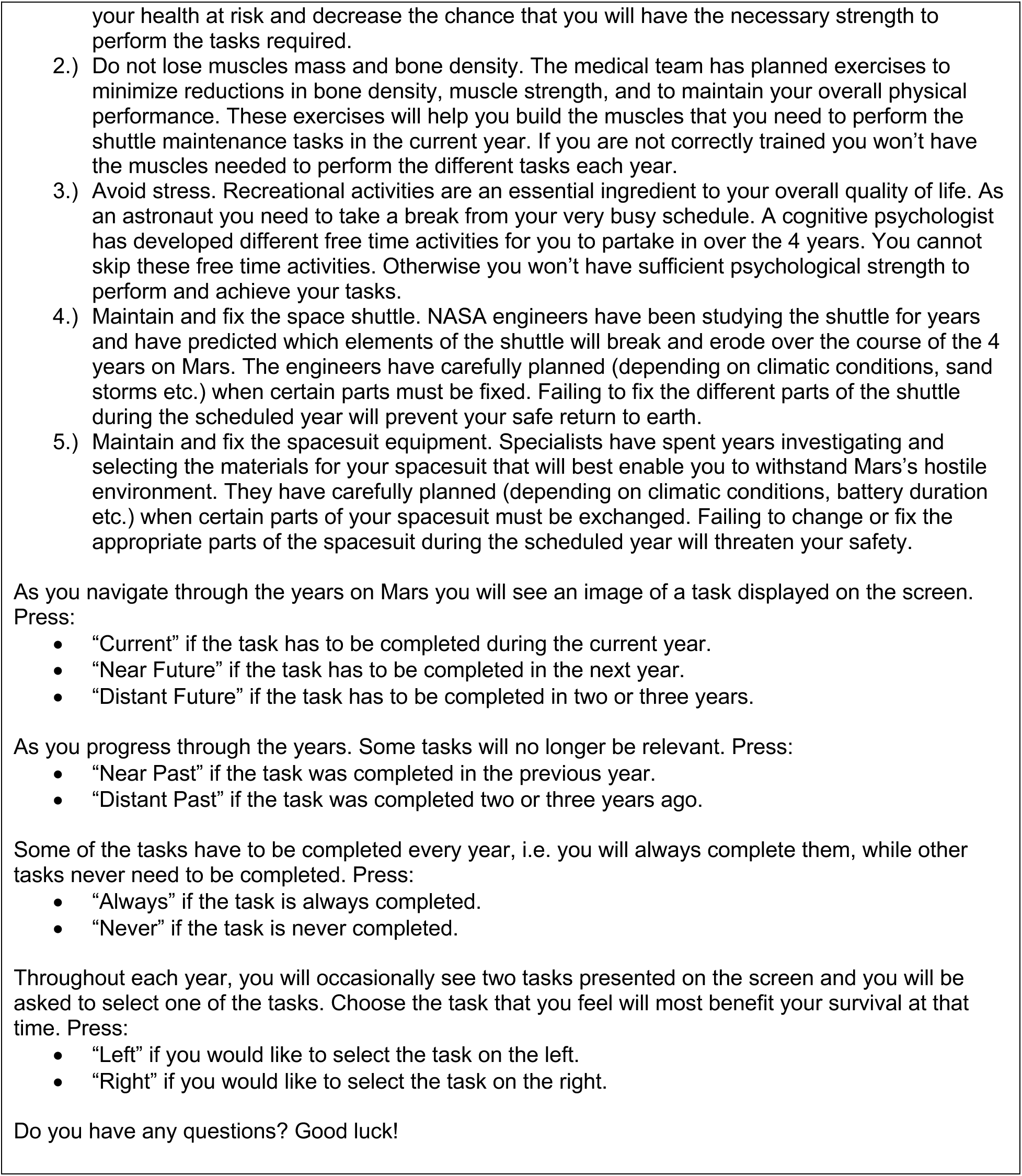
Instructions.

**Supplementary Table 7.** Temporal trajectories. The same goals were presented each year, but their temporal distance changed as participants advanced through the game. The trajectories of the goals depended on the year in which they needed to be accomplished. For instance, goals that needed to be accomplished in year 3 started off as goals relevant in the distant future, transitioned to goals relevant in the near future when the participant moved to the 2^nd^ year of their Mars mission, became presently relevant in the 3^rd^ year, and transitioned to something participants accomplished in the near past in their 4^th^ year. We chose this design, rather than employing naturalistic goals from participants’ personal pasts and futures, to isolate temporal distance. By rotating the temporal distance of the same stimuli, this design sought to ensure that our findings were related to the temporal distance of the goals and could not be explained by differences in experiential properties between past, future, close, or far constructions.

|  | Mars year 1 ----- Mars year 2 ----- Mars year 3 ----- Mars year 4 |
| --- | --- |
| Goals should be accomplished in<br>**year 1** | Current ----- Near past ----- Distant past ----- Distant past |
| Goals should be accomplished in<br>**year 2** | Near future ----- Current ----- Near past ----- Distant past |
| Goals should be accomplished in<br>**year 3** | Distant future ----- Near future ----- Current ----- Near past |
| Goals should be accomplished in<br>**year 4** | Distant future ----- Distant future ----- Near future ----- Current |

**Supplementary Table 8.** Game versions. Participants learned sets of 5 goals that needed to be accomplished in year 1, year 2, year, 3, year 4, always, and never. We created six versions of the task to counterbalance all sets of stimuli. In version 1 of the task, participants needed to accomplish a given set of goals in year 1, in version 2 of the tasks participants needed to accomplish those same goals in year 2, in version 3 they needed to be accomplished in year 3, and so on. This rotation method ensured that each set of goals was assigned to every category and that any differences observed during the task could not be attributed to a stimuli-year combination being more memorable than the others.

| Version | Number of Participants |
| --- | --- |
| V1 | 6 |
| V2 | 6 |
| V3 | 5 |
| V4 | 5 |
| V5 | 4 |
| V6 | 5 |

**Supplementary Table 9.** GLM regressors for the primary analysis. Regressors included in the whole brain and hippocampal GLMs. These GLMs examined differences in neural activity between the temporally removed goals and the current goals. All trials were modeled with boxcars, where the duration was set to the participant’s reaction time, and convolved with FEAT’s gamma hemodynamic response function. Temporal derivatives and temporal filtering was applied to regressors 1-8. Motion parameter files represented the translation and rotation values outputted from the ME-ICA preprocessing pipeline.

| ID | Regressors | Explanation |
| --- | --- | --- |
| 1-4 | remote, current, always, never | Regressors denoting the timing for all correct trials in each temporal condition. Temporally removed conditions contained a maximum of 15 trials each, while current, always, and never conditions contained a maximum of 20 trials each. |
| 5-8 | year 1, year 2, year 3, year 4 | Regressors denoting the timing for all trials in each game year, including: the task, additional choice, incorrect response, and reaction time outlier trials. |
| 9-14 | mp1, mp2, mp3, mp4, mp5, mp6 | Six motion parameters (mp) capturing the translations and rotations. |

**Supplementary Table 10.**
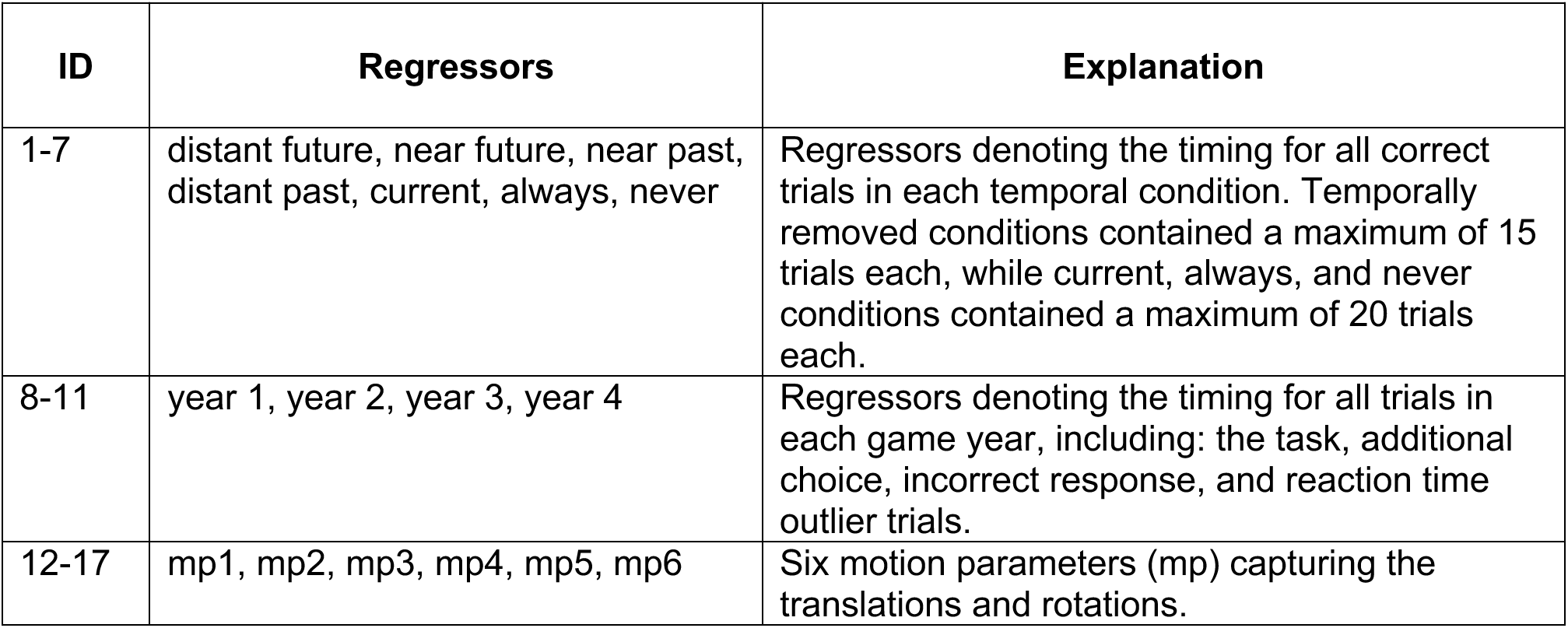
GLM regressors for the secondary analysis. Regressors included in the whole brain and hippocampal GLMs. These GLMs examined differences in neural activity between the temporally removed goals and the current goals. All trials were modeled with boxcars, where the duration was set to the participant’s reaction time, and convolved with FEAT’s gamma hemodynamic response function. Temporal derivatives and temporal filtering was applied to regressors 1-11. Motion parameter files represented the translation and rotation values outputted from the ME-ICA preprocessing pipeline.

